# PI(4)P recruits CIDE proteins to promote the formation of unilocular lipid droplets during adipogenesis and hepatic steatosis

**DOI:** 10.1101/2024.08.22.609036

**Authors:** Jin Wu, Mingming Gao, Xiaoqin Wu, Yang Liu, Ximing Du, Yan Liang, Chengxin Ma, Peng Li, Fung-Jung Chen, Hongyuan Yang

## Abstract

Lipid droplets (LDs) are evolutionarily conserved organelles that play important roles in metabolism. Each LD is enclosed by a monolayer of phospholipids, distinct from bilayer membranes. The composition of LD surface phospholipids and their impact on LD growth and function remain to be clearly defined. Phosphoinositides mark cellular organelles and regulate cell signalling and organellar function. Here, we demonstrate that PI(4)P decorates a subset of LDs to recruit and activate the CIDE proteins. Enhanced expression of ORP2 and ORP5, LD-associated lipid transfer proteins that remove PI(4)P from LDs, abolished the localization and function of CIDE proteins. Blocking the synthesis of PI(4)P on LD surface via knocking down PI4K2A also impaired the localization and function of CIDE proteins. In adipocytes, depleting PI(4)P dramatically reduced the size of LDs, as well as adipose tissue mass. In severe steatotic liver, depleting PI(4)P reduced LD accumulation. Our results thus identify a key function of LD surface PI(4)P under physiological conditions and unveil how CIDE proteins are recruited to LDs.

## INTRODUCTION

Lipid droplets (LDs) are intracellular organelles that store lipids and regulate energy homeostasis^1–6^. LDs are also actively involved in other cellular processes such as gene expression, intracellular lipid and membrane trafficking, viral replication and inflammation. Each LD contains a hydrophobic neutral lipid core of triacylglycerols (TAGs) and cholesteryl esters (CEs), enclosed by a monolayer of amphipathic lipids with proteins embedded. The composition of LD surface lipids is an important factor for the targeting of LD-associated proteins, and for LD growth and degradation^5, 7^.

The cell death-inducing DFFA-like effector (CIDE) proteins are important LD-associated proteins that promote LD growth by mediating the transfer of neutral lipids from small to large LDs^8–10^. CIDE deficiency results in the accumulation of small LDs in various cell types in both mice and humans^11^. CIDE proteins show tissue specificity: white adipocytes express abundant CIDEC; brown/beige adipocytes CIDEA/C and hepatocytes CIDEB. Notably, CIDEC is significantly upregulated during hepatic steatosis^12^. CIDE proteins play important roles in human biology and disease. For instance, CIDEA/C is the determining factor of adipocyte uni/multilocularity^13^. Loss of CIDEB function confers substantial protection against human liver diseases^14^. Despite the critical roles of CIDE proteins in LD biology and human physiology, exactly how they are recruited to LD surface remains to be elucidated.

Phosphoinositides mark cellular organelles and regulate cell signalling and organellar function. Oxysterol binding protein (OSBP) and its related proteins (ORP, for <u>O</u>SBP related <u>p</u>rotein) family of lipid transfer proteins (LTPs) have now been established as major regulators of cellular phosphatidylinositol (PI) 4-phosphate/PI(4)P metabolism^15–19^. There are 12 OSBP/ORP members in humans and 7 homologous proteins (Osh1-7) in the budding yeast *Saccharomyces cerevisiae*^20^. These proteins all share a conserved <u>O</u>SBP related <u>d</u>omain (ORD) of ∼400 amino acids located at the C-terminus of OSBP and known to bind and transfer lipids. Most members of the ORP family also possess a Pleckstrin homology (PH) domain and an FFAT (diphenylalanine in an acidic tract) motif for membrane targeting. Through the PH domain and FFAT motif, some ORPs can simultaneously bind two membranes, promoting the formation of membrane contact sites^21^. Importantly, ORPs must consume PI(4)P to transfer other lipids such as sterols and phosphatidylserine between organelles at membrane contact sites^22–25^. For instance, OSBP consumes about half of the total cellular pool of PI(4)P and is a major determinant of Golgi PI(4)P homeostasis^26^. Among the OSBP/ORPs, only ORP2 and ORP5 are known to associate with LDs^27^.

Few studies have examined the existence of phosphoinositides on LD surface, and much less is known about their function(s) at LD surface. We previously detected PI(4)P on LD surface in ORP5-deficient, but not wild type HeLa cells^27^. Here, we show that PI(4)P decorates a subset of LDs to recruit and activate the CIDE family proteins. Most importantly, we identify abundant PI(4)P on LD surface of differentiating adipocytes and uncover critical roles of PI(4)P in adipose tissue homeostasis and hepatic steatosis.

## RESULTS

### ORP2/5 and CIDEB/C localize to distinct populations of LDs

While studying the subcellular distribution of ORP2 by examining the colocalization between ORP2 and several well-known LD-resident proteins, we noticed a striking distribution pattern: ORP2 and CIDEC appear to repel each other and localize to distinct populations of LDs in both 3T3L1 and HeLa cells (Fig. 1a). By contrast, ORP2 and other well-known LD-associated proteins showed strong colocalization (Fig. S1a). A liver-specific member of the CIDE family, CIDEB, and ORP2 also localize to distinct populations of LDs in Huh 7 cells (Fig. S1b). Besides ORP2, ORP5 is the only other member of the ORP family known to exist on LDs^27^. As with ORP2, ORP5 and CIDEC also localizes to distinct populations of LDs (Fig. 1b). Together, these data demonstrate that LD-associated ORP proteins cannot co-exist with CIDE proteins on LDs (Fig. 1a-1c).

**Figure 1.**
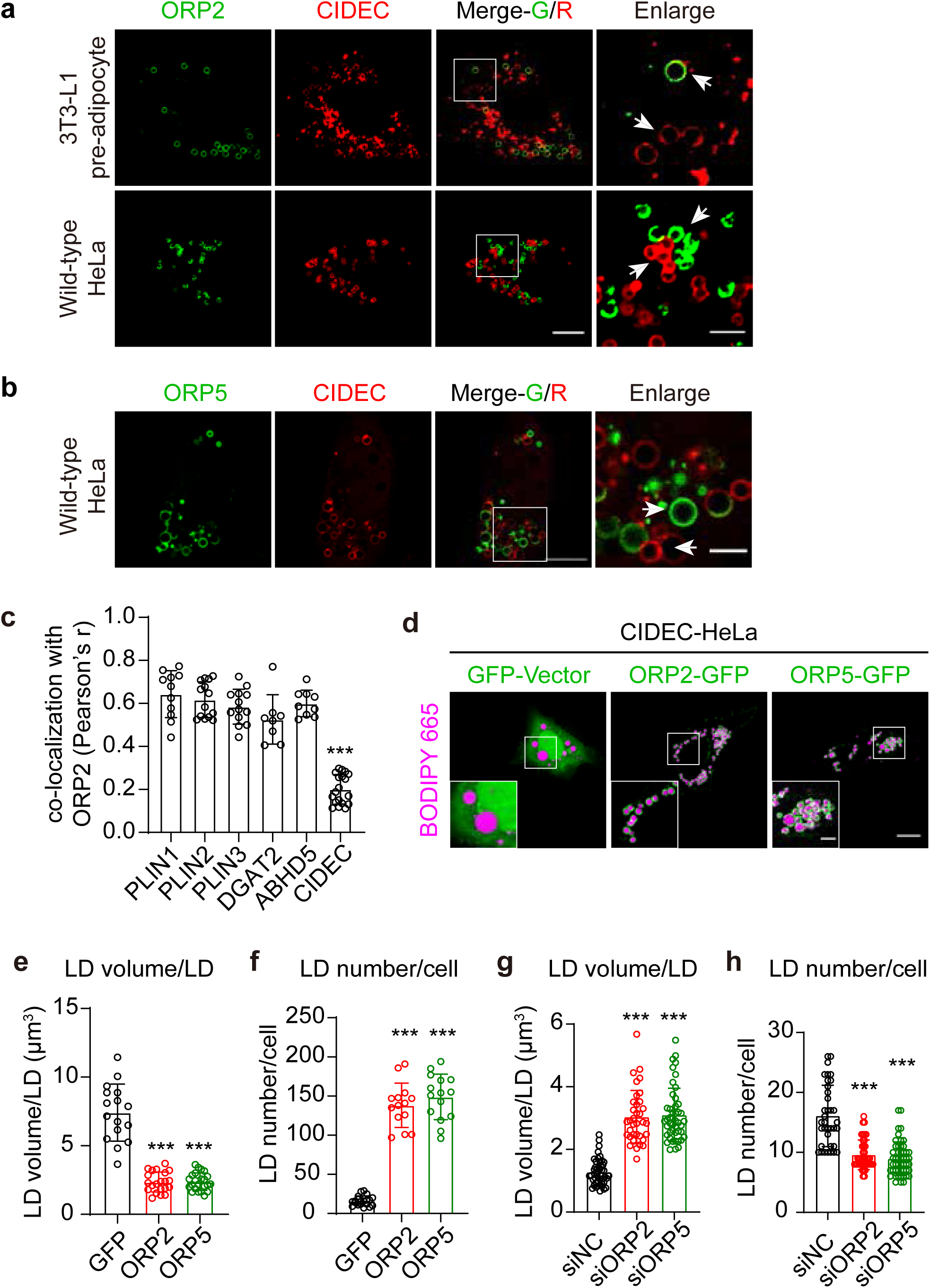
ORP2/5 impairs the LD localization of CIDE proteins and ORP2/5 inhibits CIDEC-mediated LD growth. a. Representative confocal images showing the subcellular localization of ORP2 and CIDEC in 3T3-L1 pre-adipocytes (upper) and wild-type HeLa cells (lower) treated with 200 μM OA for 16 h. Red, CIDEC. Green, ORP2. Scale bars, 10 µm; (Enlarged) 3 µm. b. Representative confocal images showing the localization of ORP5 and CIDEC in wild-type HeLa cells treated with 200 μM OA for 16 h. Red, CIDEC. Green, ORP5. Scale bars, 10 µm; (Enlarged) 3 µm. c. Quantification of the degree of co-localization between indicated proteins and CIDEC in Fig 1a, 1b and S1a. (n=8-15) The degree of colocalization is indicated by Pearson’s correlation coefficient. d. Representative confocal images showing the subcellular localization of ORP2 or ORP5 and LD morphology in HeLa cells stably expressing CIDEC (CIDEC-HeLa). Green, GFP, GFP-ORP2, and GFP-ORP5. Magenta, LDs (BODIPY 665). Scale bars, 10 µm; (Enlarged) 3 µm. e and f. Histograms showing the quantification of LD volume/LD (e) and LD number/cell (f) in CIDEC-HeLa cells overexpressing ORP2 or ORP5 after treatment with 200 μM OA for 16 h in (D). (n=15-25) g and h. Histograms showing the quantification of LD volume/LD (g) and LD number/cell (h) in CIDEC-HeLa cells after *ORP2* or *ORP5* knockdown after treatment with 50 μM OA for 16 h. (n=40-50)

### ORP2/5 inhibits CIDEC-mediated LD growth

The fact that ORP2/5 can disrupt the LD localization of CIDEB/C suggests that ORP2/5 may compromise CIDEB/C function. To examine this possibility, we employed HeLa cells stably overexpressing CIDEC (CIDEC-HeLa) where the uniquely enlarged LDs are formed by CIDEC^13, 28^. Upon expression, ORP2 or ORP5 significantly reduced the size of LDs and increased their number in CIDEC-HeLa cells (Fig. 1d-1f). This effect is specific to ORP2/5 because overexpressing other LD associated proteins (PLIN1-3 and RAB18) did not reduce LD size in these cells (Fig. S1c-S1d). Moreover, overexpressing other members of ORP family that cannot associate with LDs, did not impact CIDEC function (Fig. S1e). Knocking down ORP2/5 from CIDEC-HeLa cells increased LD size and reduced LD number (Fig. 1g-1h; S1f-S1g). Importantly, knocking down other members of ORP family, ORP1, ORP4, ORP9 or OSBP did not affect the size of LDs in CIDEC-HeLa cells (Fig. S2a-S2e). Together, these data demonstrate that ORP2/5 can specifically interrupt CIDEC-mediated LD growth.

### LD targeting is critical for ORP2/5 to impact CIDE function

Since the non-LD targeting members of ORP family had little effect on the function of CIDEC (Fig. S1e and S2e), we wonder if LD localization is a prerequisite for ORP2/5 to disrupt CIDE localization and function. We first treated cells with 22(R)-OHC (22R-hydroxycholesterol), a lipid known to trap ORP2 at the PM^29^. Indeed, 22(R)-OHC abolished the LD localization of ORP2 and accordingly, increased LD size in CIDE-C HeLa cells (Fig. S2f-S2g). We next investigated the LD targeting domains/motifs of ORP2/5. The FFAT motif of ORP2 and the transmembrane domain of ORP5 are known to be required for their localization to the ER (Fig. 2a)^27, 29^. Removing the FFAT motif of ORP2 or the transmembrane region of ORP5 also abolished their LD localization, suggesting the ER localization of ORP2 and ORP5 is essential to their LD association (Fig. 2b and 2e). Both ORP2 and ORP5 also possess an amphipathic helical lid region before the lipid binding/transferring ORD domain (Fig. 2a and 2c)^29^. The helical regions contain putative large, LD-targeting hydrophobic residues^30^, e.g. W77 and I79 of ORP2, and W372 and L374 of ORP5 (Fig. 2a). Removing the helical regions or mutating the W, I or L residues significantly impaired the LD targeting of ORP2 and ORP5 (Fig. 2d-2e). Importantly, all ORP2 and ORP5 mutants with defective LD targeting had little effect on LD growth in CIDEC HeLa cells (Fig. 2f-2g). Finally, the same mutants also failed to impact LD growth in CIDEA-HeLa cells (Fig. S3a-S3b). Together, these data identify LD targeting mechanisms of ORP2/5 and demonstrate that ORP2/5 must localize to LDs to impede the function of CIDE proteins.

**Figure 2.**
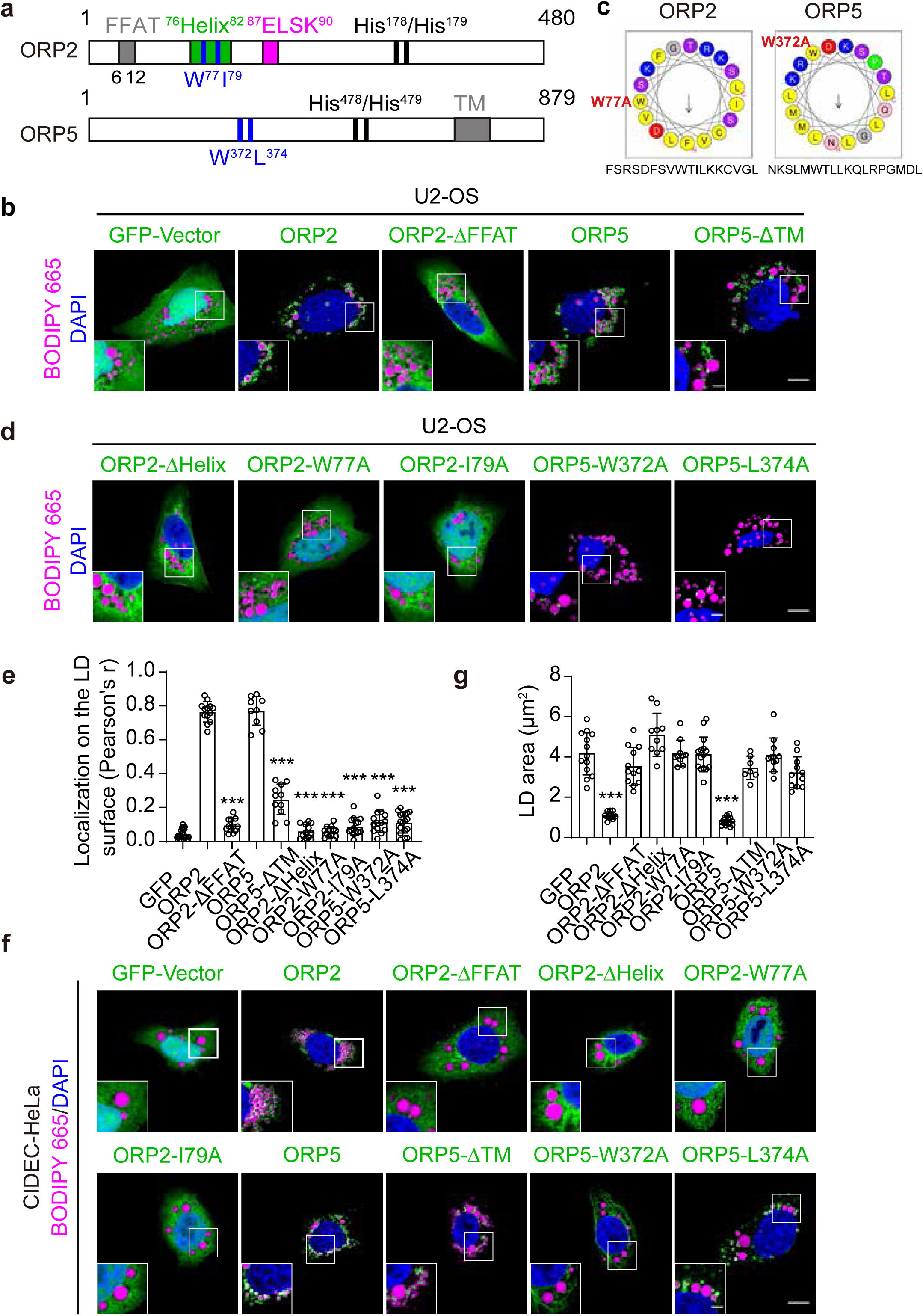
Amino acids with large hydrophobic side group in ORP2/5 determine the LD localization of ORP2/5 and their impact on CIDEC. a. Schematic diagram showing the amino acid sequences of ORP2 and ORP5. b. Representative confocal images showing the localization of GFP-tagged ORP2, ORP2-ΔFFAT, ORP5, and ORP5-ΔTM in U2-OS cells. Green, GFP and GFP-tagged indicated proteins. Magenta, LDs (BODIPY 665). Blue, nuclei (DAPI). Scale bars, 10 µm; (Enlarged) 3 µm. c. Schematic diagram showing the amphipathic α-helical region within ORP2 and ORP5. d. Representative confocal images showing the subcellular localization of GFP-tagged ORP2-ΔHelix, ORP2-W77A, ORP2-I79A, ORP5-W372A and ORP5-L374A in U2-OS cells. Green, GFP and GFP-tagged indicated proteins. Magenta, LDs (BODIPY 665). Blue, nuclei (DAPI). Scale bars, 10 µm; (Enlarged) 3 µm. e. Histogram showing the quantification of LD localization of GFP-tagged proteins in (b) and (d). (n=8-15) LD indicated by overexpression of PLIN1-RFP, LD localization of ORPs and ORPs-mutant is represented by Pearson’s correlation coefficient of two fluorescence signals. f. Representative confocal images showing the subcellular localization of GFP-tagged ORP2, ORP2-ΔFFAT, ORP2-ΔHelix, ORP2-W77A, ORP2-I79A, ORP5, ORP5-ΔTM, ORP5-W372A, and ORP5-L374A and LD morphology in CIDEC-HeLa cells treated with 200 μM OA for 16 h. Green, GFP and GFP-tagged indicated proteins. Magenta, LDs (BODIPY 665). Blue, nuclei (DAPI). Scale bars, 10 µm; (Enlarged) 3 µm. g. Histogram showing the quantification of average LD area regulated by the expression of indicated proteins in (f). (n=8-15)

### ORP2/5 impairs LD localization of CIDEC and inhibits CIDEC-mediated lipid exchange

CIDE proteins mediate LD fusion/growth in three steps: 1. Localization of CIDE proteins to LDs; 2. Enrichment/condensation of CIDE proteins at LD-LD contact sites; 3. Formation of lipid permeable passageways for neutral lipids to flow from small to large LDs^31^. We wonder which step(s) is affected by LD-localized ORP2/5. We first generated CIDEC-HeLa cells stably expressing ORP2/5 and purified LDs from these cells. The amount of CIDEC, but not PLIN3 or ABHD5, was much reduced from the LD fraction upon ORP2/5 overexpression (Fig. 3a). Total cellular level of CIDEC was not affected by ORP2/5 overexpression (Fig. 3a). CIDEC forms condensates at LD-LD contact sites, and ORP2/5 also significantly reduced the formation of these condensates (Fig. 3b). We further examined the rate of lipid exchange between a large number of random LD pairs as we described previousy^9, 13, 31^. ORP2/5 expression dramatically reduced CIDEC-mediated lipid exchange between LDs (Fig. 3c-3g).

**Figure 3.**
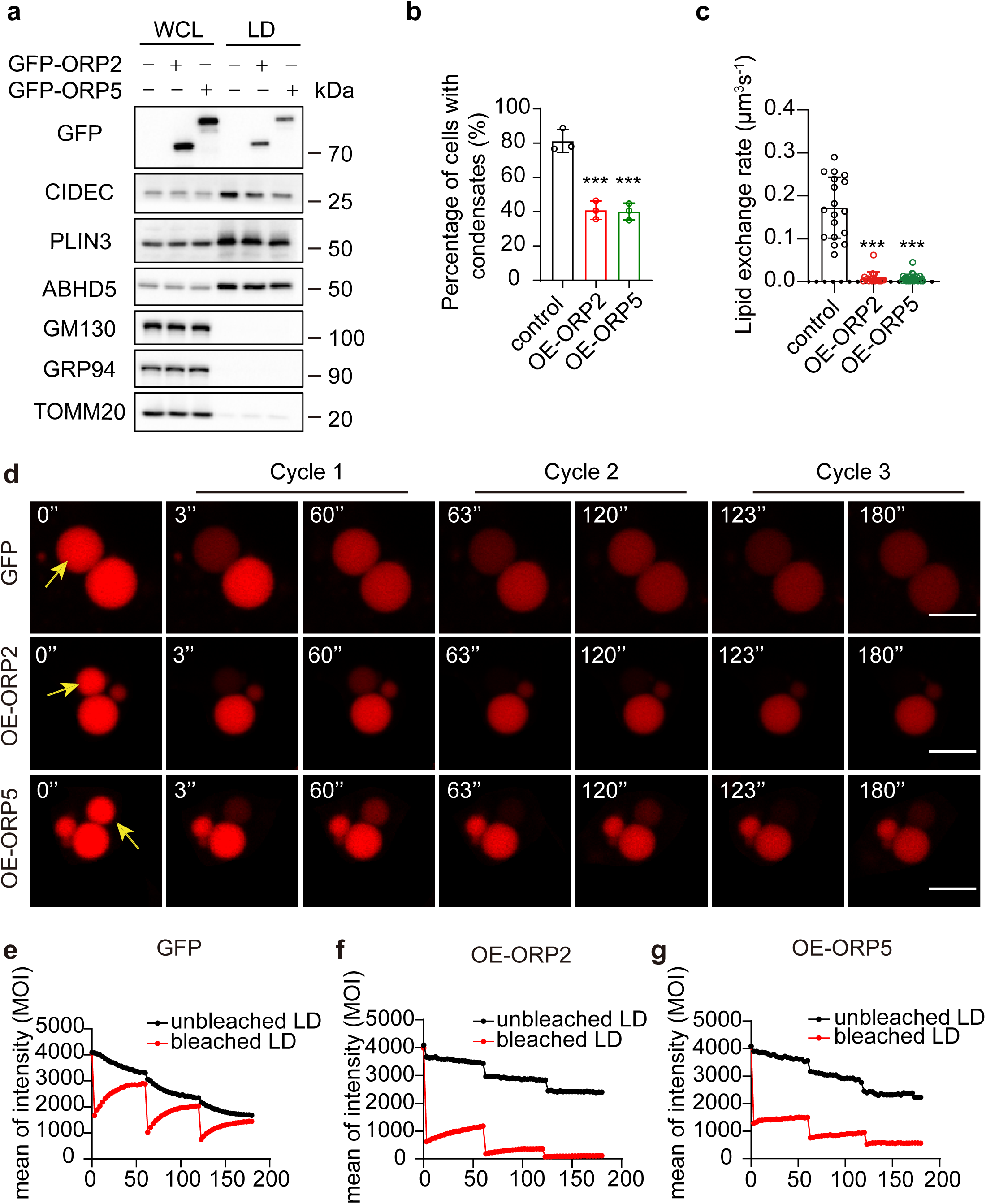
ORP2/5 impair the LD localization of CIDEC and CIDEC condensation at LD-LD contact to inhibit CIDEC-mediated lipid exchange and LD fusion. a. Western blotting showing the level of CIDEC and ORP2/5 proteins in whole cell lysate (WCL) and the subcellular LD fractionation extracted from CIDEC-HeLa cells overexpressing GFP-ORP2 or GFP-ORP5. b. Histogram showing the percentage of HeLa cells with CIDEC condensates. The cells overexpressed empty vector (control), ORP2 (OE-ORP2) or ORP5 (OE-ORP5). Three independent experiments. c. Histogram showing the calculated lipid exchange rates in CIDEC-HeLa cells overexpressing ORP2 or ORP5. The measurements were performed by FRAP-based lipid exchange rate assay. (n=18-30) d-g. Representative images (d) and mean optical intensity traces (e)-(g) of FRAP-based lipid exchange rate assay for one pair of LDs in CIDEC-HeLa cells expressing GFP, GFP-ORP2, and GFP-ORP5, respectively. LDs were pre-stained with BODIPY C12. Scale bars, 3 µm.

### ORP2/5 regulates CIDE proteins through PI(4)P

We next sought to further determine how exactly the LD-localized ORP2/5 may impact CIDE protein function. Recently, CLSTN3β was reported to inhibit CIDE protein function likely through direct interaction^13^. However, we were not able to detect any interaction between CIDE proteins and ORP2 (Fig. S3c), suggesting that ORP2 may regulate CIDE proteins indirectly. Since ORP2 and ORP5 are lipid transfer proteins, we wonder if ORP2/5 may regulate CIDE function through LD surface lipids. ORP2 and ORP5 deliver different lipids (cholesterol for ORP2 and phosphatidylserine for ORP5) to LDs, but they both must consume PI(4)P on LD surface during the delivery process^27^. Indeed, cells expressing wild type ORP2/5 showed reduced PI(4)P signal on LD surface (Fig. 4a-4b). An ORP2 mutant without its cholesterol binding motif (ELSK) and therefore defective in delivering cholesterol can still inhibit CIDEC-mediated LD growth (Fig. 2a, 4c-4d). Likewise, ORP5 with compromised phosphatidylserine binding (L389D) also retained its ability to inhibit LD growth (Fig. 2a, 4c-4d). By contrast, changing the highly conserved histidine residues (critical for PI(4)P binding and transfer) of ORP2/5 to alanine abolished their inhibitory effects on LD growth in CIDEC HeLa cells (Fig. 2a, 4c-4d).

**Figure 4.**
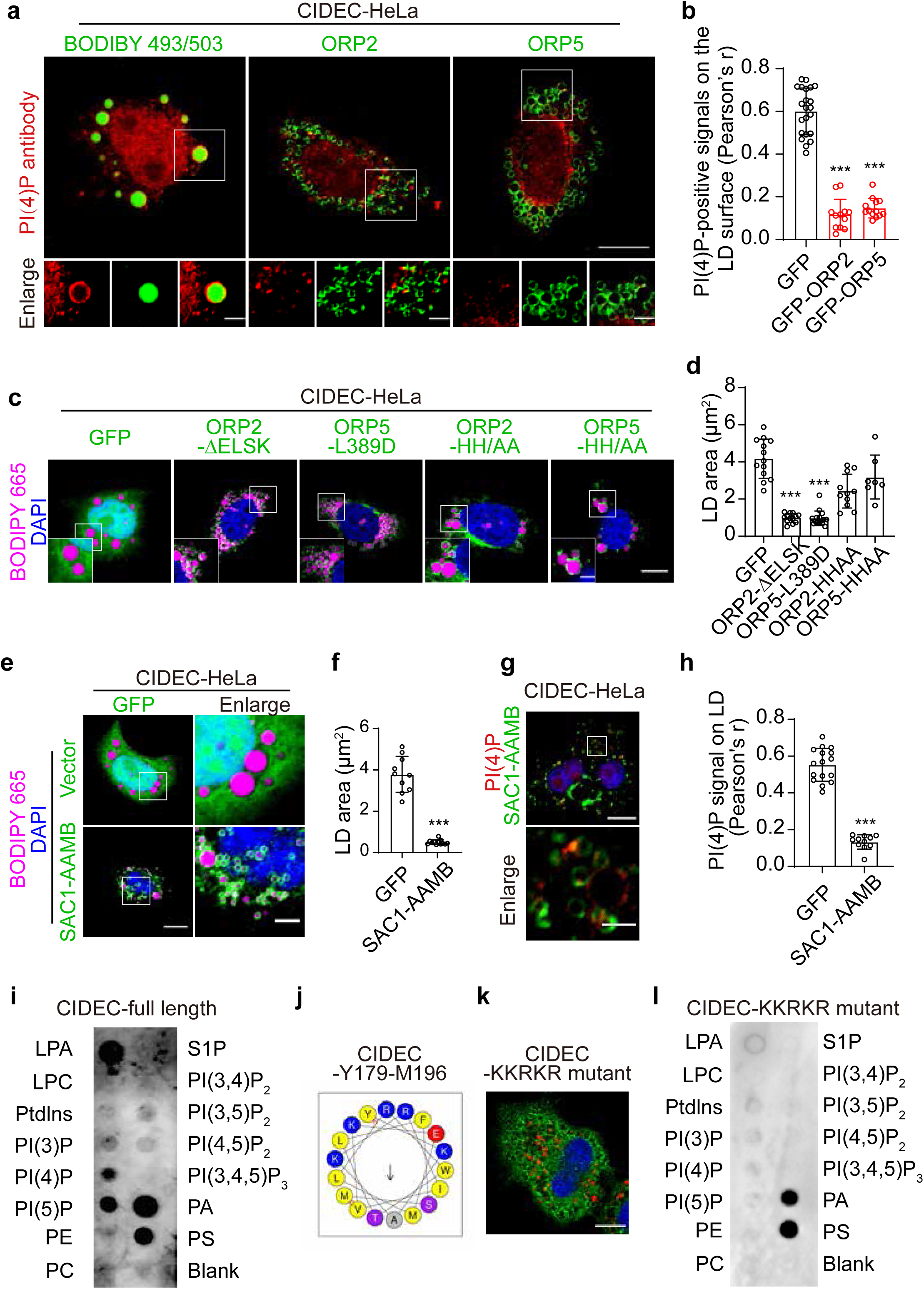
ORP2/5 reduces the level of PI(4)P on LDs to inhibit CIDEC-mediated LD growth. a. Representative confocal images showing PI(4)P-positive signals on the LD surface detected by PI(4)P antibody in CIDEC-HeLa cells overexpressing GFP-ORP2 or GFP-ORP5. Green, LDs (BODIPY 493/503), ORP2, or ORP5. Red, PI(4)P. Scale bars, 10 µm; (Enlarged) 3 µm. b. Histogram showing the quantification of PI(4)P-positive signals on the LD surface in (a). (n=12-22) c. Representative confocal images showing the localizations of ORP2-ΔELSK (binding with cholesterol), ORP5-L389D (binding with PS), ORP2-HH/AA (binding with PI(4)P), ORP5-HH/AA (binding with PI(4)P) in CIDEC-HeLa cells treated with 200 μM OA for 16 Hrs. Green, GFP and ORP2 or ORP5 mutants. Magenta, LDs (BODIPY 665). Blue, nuclei (DAPI). Scale bars, 10 µm; (Enlarged) 3 µm. d. Histogram showing the quantification of average LD area in (c). (n=7-15) e. Representative confocal images showing LD morphology in CIDEC-HeLa cells overexpressing SAC1-AAMB (AAMB, LD targeting sequence). Green, GFP and SAC1-AAMB. Magenta, LDs (BODIPY 665). Blue, nuclei (DAPI). Scale bars, 10 µm; (Enlarged) 3 µm. f. Histogram showing the quantification of LD area in (e). (n=10-15) g. Representative confocal images showing PI(4)P-positive signals in CIDEC-HeLa cells overexpressing SAC1-AAMB-GFP. Scale bars, 10 µm; (Enlarged) 3 µm. h. Histogram showing the quantification of PI(4)P-positive signals on LD in CIDEC-HeLa cells overexpressing SAC1-AAMB-GFP compared to GFP vector for Fig 4g. (n=11-16) i. Lipid strip assay showing the affinity of purified full length CIDEC for phospholipids. j. Schematic diagram showing the amphipathic feature of the α-helix within CIDEC-Y179-M196. k. Confocal images showing the localizations of CIDEC-KKRKR mutant in HeLa cell. Green, CIDEC-KKRKR mutant. Red, LDs (BODIPYC12). Blue, nuclei (DAPI). Scale bars, 10 µm. l. Lipid strip assay showing the affinity of CIDEC KKRKR mutant for phospholipids.

These data suggest that PI(4)P, but not cholesterol nor phosphatidylserine, may directly impact CIDE function. If so, then reducing LD surface PI(4)P by other means should also inhibit CIDE function. Since the non-LD targeting members of ORP family can also consume PI(4)P, we attached the LD targeting motif of AAM-B to OSBP, ORP1S, ORP4S and ORP9S^32^. These fusion proteins can now localize to LD surface, and importantly, reduce LD size in CIDEC-HeLa cells (Fig. S4a-S4b). To further examine the role of LD surface PI(4)P in CIDE function, we fused the LD-targeting motif of AAM-B to SAC1, a PI(4)P phosphatase. SAC1-AAMB localized to LD surface, reduced LD surface PI(4)P and LD size in CIDEC-HeLa cells (Fig. 4e-4h). Finally, purified CIDEC can directly interact with PI(4)P and other anionic phospholipids *in vitro*, suggesting LD surface PI(4)P may directly interact with and recruit CIDEC (Fig. 4i). CIDEC possesses a predicted amphipathic helical region between a.a. Y179 and T217 (Fig. 4j and S4c), which is essential to LD targeting and PI(4)P binding of CIDEC (Fig. S4d-S4e). Replacing the positively charged residues within this helical region with alanine (KKRKR mutant) also abolished PI(4)P binding and LD targeting of CIDEC (Fig. 4k-4l). While CIDEC can also bind PI5P in the lipid-strip assay (Fig. 4i), we were not able to detect any PI5P on LDs of CIDEC HeLa cells (data not shown). Together, these data strongly indicate that LD surface PI(4)P can directly regulate the recruitment and function of CIDE proteins.

### PI(4)P produced by PI4K2A promotes CIDE protein function

PI(4)P is synthesized by PI-4 kinases, therefore, we next determined if any PI-4 kinase is critical for CIDE function. Knocking down PI4K2A, but not the other three PI-4 kinases reduced LD size and increased LD number in CIDEC-HeLa cells (Fig. 5a-5b, S5a-S5d). Re-expressing siRNA resistant forms of WT PI4K2A, but not the catalytic dead mutants, restored LD size in PI4K2A-deficient CIDEC HeLa cells (Fig. 5c and 5d). As with overexpressing ORP2/5 (Fig. 3), knocking down PI4K2A also reduced LD localization of CIDEC, the enrichment of CIDEC at LD-LD contact sites and the rates of lipid exchange between random LD pairs (Fig. 5e-5g; Fig. S5e-S5g). Since ORP2/5 reduces whereas PI4K2A increases LD surface PI(4)P, we wonder if PI4K2A can neutralize the effect of ORP2/5. Expressing ORP2/5 reduced LD size of CIDEC-HeLa cells, but co-expressing a LD-specific PI4K2A (PI4K2A-AAMB) reversed these changes (Fig. 5h and 5i). These data again reinforced a critical role for PI(4)P in controlling the localization and function of CIDE proteins.

**Figure 5.**
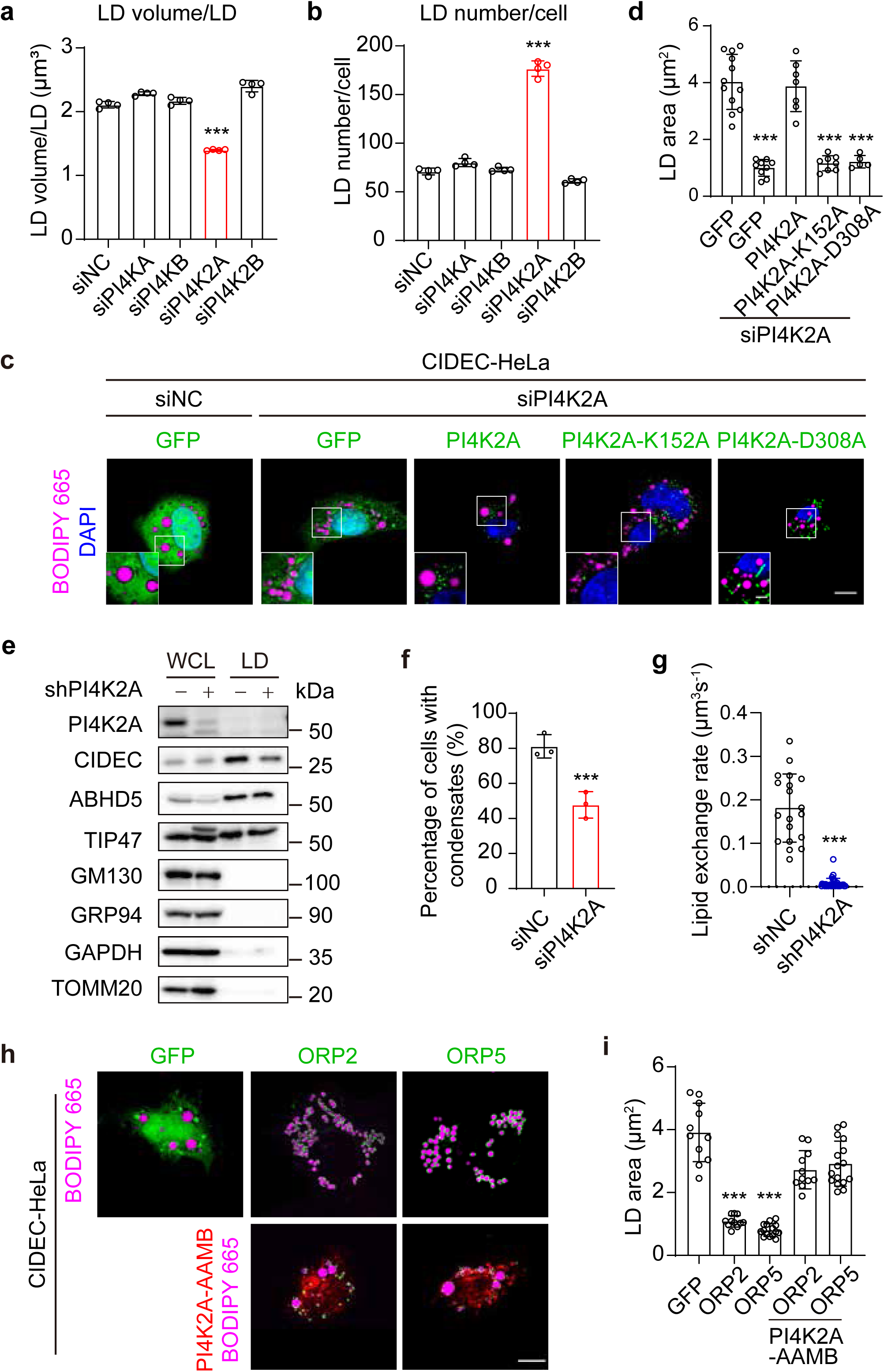
PI(4)P synthesized by PI4K2A promotes the LD localization of CIDE proteins and their condensation at LD-LD contact. a and b. Histograms showing the quantification of LD volume/LD (a) and LD number/cell (b) in CIDEC-HeLa cells after treatment with scramble siRNA (siNC) or four indicated siRNAs (siPI4KA, siPI4KB, siPI4K2A, and siPI4K2B) analyzed by high-content screening system for four independent experiments. c. Representative confocal images of LDs in CIDEC-HeLa cells after treatment with scramble siRNA (siNC) or *PI4K2A* siRNA (siPI4K2A), followed by overexpressing GFP vector, wild-type PI4K2A, or kinase-dead PI4K2A mutants (PI4K2A-K152A, PI4K2A-D308A). Green, GFP, wild-type PI4K2A, and PI4K2A mutants. Magenta, LDs (BODIPY 665). Blue, nuclei (DAPI). Scale bars, 10 µm; (Enlarged) 3 µm. d. Histogram showing the quantification of average LD area in (c). (n=5-15) e. Western blotting showing the level of CIDEC protein in the subcellular LD fractionation extracted from CIDEC-HeLa cells infected with lentiviral GFP-tagged scramble shRNA (shNC) or *PI4K2A* shRNA (shPI4K2A). f. Histogram showing the percentage of wild-type HeLa cells with CIDEC condensates. The cells were co-transfected of CIDEC-GFP with scramble siRNA (siNC) or *PI4K2A* siRNA (siPI4K2A). Three independent experiments. g. Histogram showing the calculated lipid exchange rates CIDEC-HeLa cells infected with lentiviral GFP-tagged scramble shRNA (shNC) or PI4K2A shRNA (shPI4K2A). The measurements were performed by FRAP-based lipid exchange rate assay. (n=20-30) h. Representative confocal images showing LDs in CIDEC-HeLa cells co-overexpressing ORP2 or ORP5 with PI4K2A-AAMB. Green, GFP, GFP-ORP2, and GFP-ORP5. Red, PI4K2A-AAMB. Magenta, LDs (BODIPY 665). Scale bar, 10 µm. i. Histogram showing the quantification of average LD area in (h). (n=10-15)

### ORP2/5, PI4K2A and PI(4)P regulate adipocyte maturation and lipolysis

CIDEC plays a pivotal role in forming the characteristic unilocular LDs of mature white adipocytes^33, 34^. To understand the physiological function of LD surface PI(4)P, we examined PI(4)P during the differentiation of 3T3-L1 preadipocytes. Distinct from cultured cancer cell lines where PI(4)P can hardly be observed on LD surface^27^, we detected strong PI(4)P signal on day 3 of the differentiation process (Fig. 6a). Notably, PI(4)P signal decreased significantly on day 4, accompanied by a dramatic increase of LD size (Fig. 6b-6c). Most of the PI(4)P appears to associate with relatively small and growing LDs (Fig. 6d), consistent with our conclusion that PI(4)P functions to recruit CIDE proteins to promote LD growth. Moreover, PI(4)P and CIDEC also co-localize more on relatively small LDs (Fig. 6e-6f). We also detected PI4K2A on LD surface in 3T3-L1 cells three days post differentiation (Fig. S6a). Knocking down PI4K2A, but not PI4KA, dramatically reduced the amount of detectable PI(4)P (Fig. S6b), suggesting LD surface PI(4)P is also synthesized by PI4K2A in 3T3 L1 preadipocytes.

**Figure 6.**
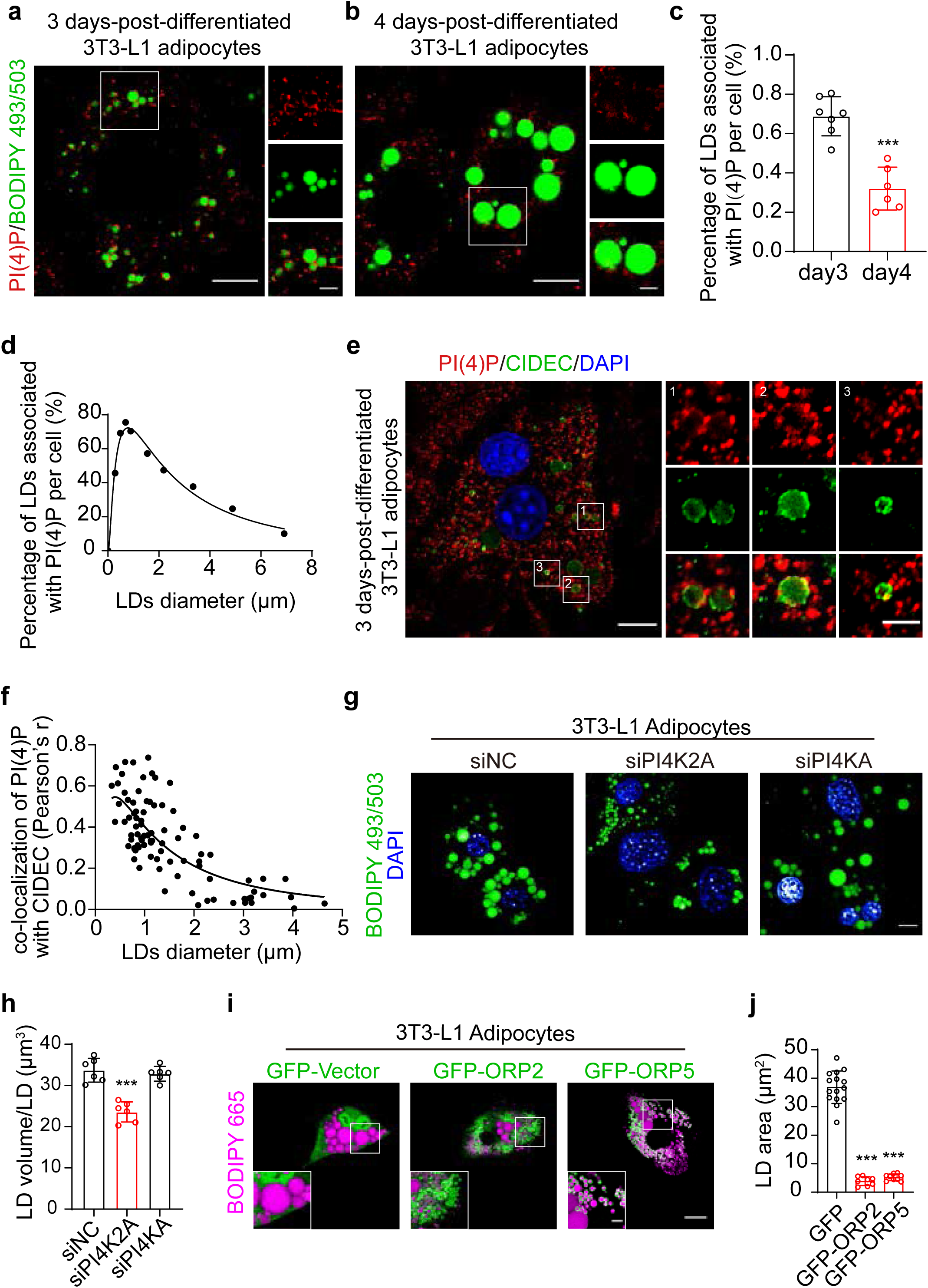
ORP2/5 and PI4K2A control adipocyte maturation through regulating the level of PI(4)P on LDs. a. Representative confocal images showing PI(4)P and LDs in 3T3-L1 adipocytes 3 days after induction of differentiation. Red, PI(4)P. Green, LDs (BODIPY 493/503). Scale bars, 10 µm; (Enlarged) 3 µm. b. Representative confocal images showing PI(4)P and LDs in 3T3-L1 adipocytes 4 days after induction of differentiation. Red, PI(4)P. Green, LDs (BODIPY 493/503). Scale bars, 10 µm; (Enlarged) 3 µm. c. The percentage of LDs associated with PI(4)P per cell in 3 in 3T3-L1 adipocytes as shown in a and b. (n=6-7) d. The percentage of LDs associated with PI(4)P per cell according to LD size in 3T3-L1 adipocytes as shown in a and b. e. Representative confocal images showing PI(4)P and the subcellular localization of endogenous CIDEC in 3 days-post-differentiated 3T3-L1 adipocytes. Red, PI(4)P. Green, CIDEC proteins. Blue, nuclei (DAPI). Scale bars, 10 µm; (Enlarged) 3 µm. f. The colocalization between PI(4)P and CIDEC according to LD size in cells as showed in e. g. Representative confocal images showing LDs in 3T3-L1 adipocyte after treatment with scramble siRNA (siNC), *PI4K2A* siRNA (siPI4K2A), or *PI4KA* siRNA (siPI4KA). Green, LDs (BODIPY 493/503). Blue, nuclei (DAPI). Scale bar, 10 µm. h. Histogram showing the quantification of LD volume/LD in 3T3-L1 adipocytes as in (g) analyzed by high-content screening system. (n=6) i. Representative confocal images showing the LDs in 3T3-L1 adipocytes overexpressing ORP2 or ORP5. Green, GFP, GFP-ORP2, and GFP-ORP5. Magenta, LDs (BODIPY 665). Scale bars, 20 µm; (Enlarged) 5 µm. j. Histogram showing the quantification of average LD area in 3T3-L1 adipocytes as in (i). (n=8-15)

We next examined the role of PI(4)P in LD dynamics during adipogenesis. Knocking down PI4K2A, but not PI4KA, PI4KB or PI4K2B reduced LD volume (Fig. 6g-6h, Fig. S6c-S6h). Similarly, overexpressing ORP2/5 in 3T3-L1 adipocytes reduced the size of LDs (Fig. 6i-6j). We further examined the physiological impact of PI(4)P loss on 3T3 L1 adipocytes under lipolytic conditions. There are decreased LD volume and increased LD number in PI4K2A-deficient 3T3 L1 adipocytes when treated with isoproterenol (Fig. S6i-S6j). Moreover, significantly more NEFA and glycerol were detected in the media of PI4K2A-deficient adipocytes treated with isoproterenol (Fig. S6k and S6l). Enhanced lipolysis is consistent with increased number and surface area of LDs in PI4K2A-deficient adipocytes.

### ORP2/5, PI4K2A and PI(4)P regulate adipose tissue homeostasis and hepatic steatosis

To further evaluate the role of LD surface PI(4)P in adipocyte function *in vivo*, we generated AAV (Adeno-associated virus) constructs overexpressing ORP5 or shRNA against PI4K2A and infected the inguinal white adipose tissue (iWAT) of mice for 2 months (Fig. 7a). Although having negligible effect on fat mass (Fig. 7b), overexpressing ORP5 or knocking down PI4K2A significantly reduced the size/area of LDs, consistent with defective CIDEC function (Fig. 7c-7d). We also carried out intraperitoneal injection of CL316243 to stimulate lipolysis after AAV infection (Fig. S7a). Overexpressing ORP2/5 or knocking down PI4K2A significantly reduced the size and weight of iWAT (Fig. S7b-S7c), as well as the LD size of adipocytes (Fig. S7d-S7e). Overexpressing ORP2/5 or knocking down PI4K2A in iWAT alone did not impact serum TAG, total plasma cholesterol and NEFA after injection of CL316243 (Fig. S7f-7h). We further induced lipolysis by fasting for 20 h after AAV infection for 2 months (Fig. 7e). Again, overexpressing ORP2/5 or knocking down PI4K2A reduced the size and weight of iWAT (Fig. 7f-7g), as well as the size of LDs (Fig. 7h-7i). Overexpressing ORP2/5 or knocking down PI4K2A in iWAT alone did not impact serum TAG, total plasma cholesterol and NEFA after fasting (Fig. S7i-S7k). Finally, CIDEC is required for developing severe liver steatosis in *ob/ob* mice^12^. We therefore overexpressed ORP2/5 and knocked down PI4K2A in *ob/ob* hepatocytes using the hepatocyte specific thyroxin binding globulin (TBG) promoter (Fig. 7j and S8a). The size/area of LDs in *ob/ob* liver was significantly reduced (Fig. 7k and S8b).

**Figure 7.**
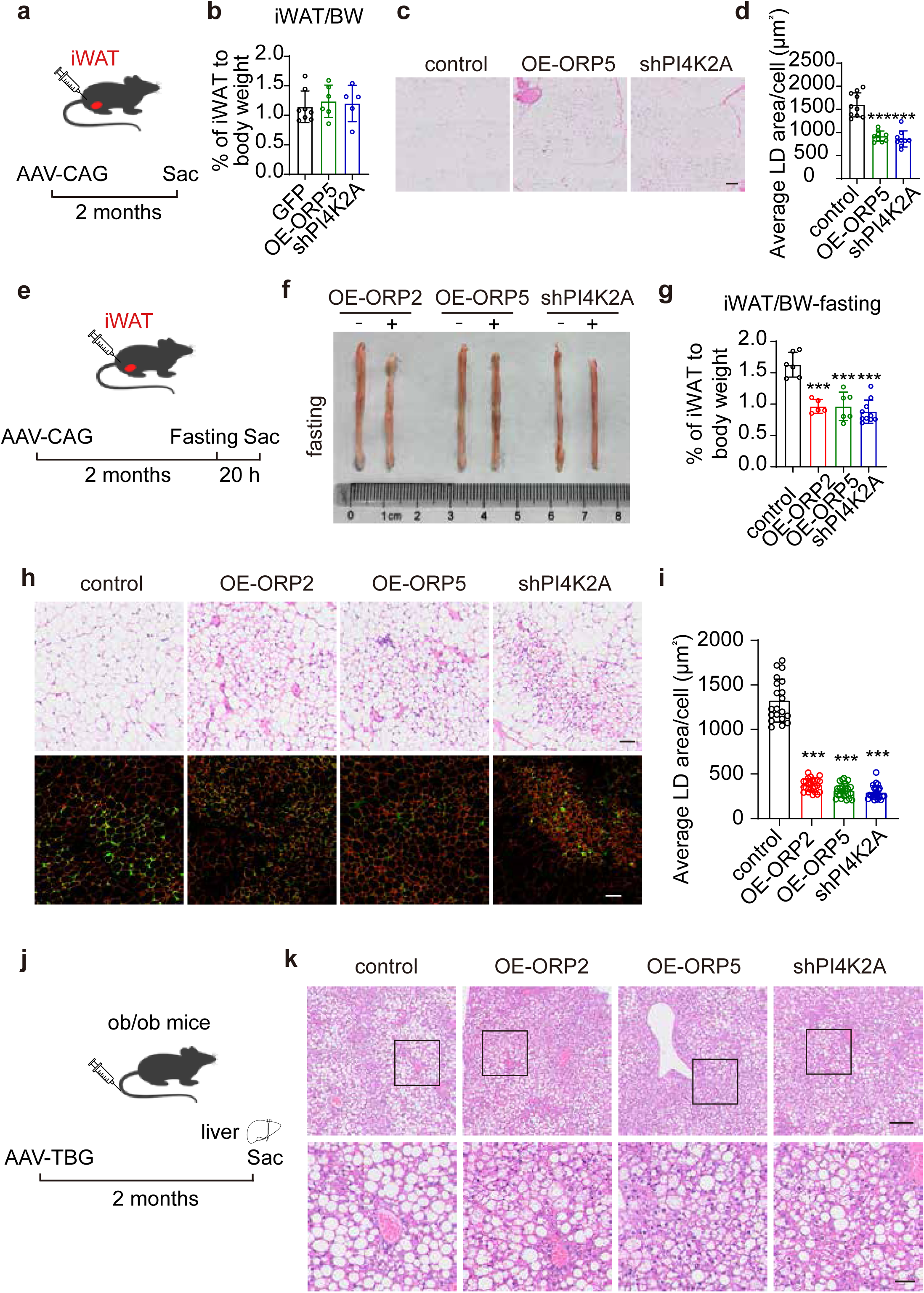
ORP2/5 overexpression or *PI4K2A* knockdown impaired LD growth in adipose tissue and in steototic liver. a. Schematic diagram showing that 8-week-old male mice were administered by *in situ* injection with AAV-CAG-GFP, AAV-CAG-GFP-ORP5, or AAV-CAG-GFP-shPI4K2A into the inguinal white adipose tissue (iWAT) for 2 months to specifically express the indicated proteins. n=5-8 mice. Sac, sacrifice. b. Histogram showing the ratio of iWAT mass to body weight (iWAT/BW) of mice from (a). c. H&E staining of the iWAT from (a). Scale bar, 100 µm. c. Histogram showing the quantification of average LD area of the iWAT in (c). e. Schematic diagram showing that 8-week-old male mice were administered by *in situ* injection with AAV-CAG-GFP, AAV-CAG-GFP-ORP2, AAV-CAG-GFP-ORP5, or AAV-CAG-GFP-shPI4K2A into the iWAT for 2 months followed by fasting for 20 h. n=5-10 mice. f. Representative images showing the iWAT of mice fasted for 20 h as in (e). g. Histogram showing the ratio of iWAT mass to body weight (iWAT/BW) of mice after fasting for 20 h. h. H&E staining and immunofluorescence images (red, Perilipin1, green, GFP) showing the iWAT of mice fasted for 20 h in (e). Scale bars, 100 µm. i. Histograms showing the quantification of average LD area of iWAT in (h). j. Schematic diagram showing that 8-week-old *ob/ob* male mice were administered by intravenous injection with AAV-TBG-GFP, AAV-TBG-Flag-ORP2, AAV-TBG-Flag-ORP5, or AAV-TBG-GFP-shPI4K2A into liver for 2 months to specifically overexpress the indicated proteins or to knock down PI4K2A. n= 5 mice. Sac, sacrifice. k. H&E staining images showing the liver morphology of *ob/ob* mice as in (j). Scale bars, 200 µm, (Enlarged) 50 µm.

## DISCUSSION

Phosphoinositides mark cellular organelles and regulate their function. Whether phosphoinositides exist on LDs, now established as a class of organelles, remains to be unequivocally determined. Moreover, the function of LD surface phosphoinositides under physiological settings has not been characterized. Here, our data convincingly demonstrate the existence of PI(4)P on LD surface in differentiating adipocytes and unveil a key physiological function of LD surface PI(4)P: recruiting CIDE proteins to promote the formation of unilocular LDs (Fig. S9). We further demonstrate that LD surface PI(4)P can regulate adipose tissue lipolysis as well as LD accumulation during hepatic steatosis.

CIDE proteins play pivotal roles in human physiology and disease^8, 11^. For instance, the formation of the characteristic unilocular LDs in white adipocytes depends on CIDEC^8, 35^. Conversely, the activity of CIDEA needs to be constrained by calsyntenin 3b (CLSTN3b) to ensure multilocularity of thermogenic adipocytes^13^. CIDEB regulates lipid storage and lipoprotein secretion in the liver, and impaired CIDEB function has been linked with lower odds of developing liver disease from any cause^14^. By concentrating at LD-LD contact sites, CIDEA/B/C mediates the transfer of neutral lipids from small to large LDs in a few steps^9, 11^. The initiating step is the recruitment of CIDE proteins to LDs. Despite intensive research efforts, exactly how CIDE proteins reach LD surface remains unclear. Here we provide a few lines of evidence that LD surface PI(4)P is necessary for the LD localization of CIDE proteins. CIDE proteins cannot reach LDs decorated by ORP2/5, lipid transfer proteins known to remove PI(4)P from LDs. PI4K2A produces PI(4)P on LD surface and its deficiency impairs LD targeting of CIDE proteins. Importantly, orthogonal approaches to remove LD surface PI(4)P, e.g., forcing SAC1 onto LD surface, also abolished the localization of CIDE proteins to LDs. We further identified positively charged residues (K180, K182, R183, K186, and R190) in the amphipathic helical region of CIDEC which are critical for its PI(4)P binding and LD targeting. Although ORP2/5 and PI4K2A can impact other cellular processes besides LD surface PI(4)P, the combined results from these studies clearly indicate the LD surface PI(4)P is a crucial factor for recruiting CIDE proteins to LDs.

Among all cellular organelles, LDs are highly unique in that each LD is bounded by a monolayer of phospholipids, which play key roles in LD budding, growth, degradation, as well as protein targeting^2, 5, 36^. Phosphoinositides, quantitatively minor phospholipids, are known to govern the identity and function of cellular organelles^37^. We previously identified PI(4)P on LD surface and defined its role in driving the delivery of phosphatidylserine to LDs through ORP5^27^. However, LD surface PI(4)P most likely serves as a signal to recruit various proteins to help establish the unique identity and function of LDs^38^. Here we unveil critical roles for PI(4)P in controlling the dynamics of LDs during adipogenesis and during the development of severe hepatic steatosis. While our data strongly suggest that these effects of PI(4)P arise from its role in recruiting CIDE proteins, we cannot rule out other possibilities. For instance, PI(4)P may also recruit/regulate other LD-associated proteins. Future studies will undoubtedly uncover additional mechanisms by which PI(4)P may contribute to LD function and lipid storage.

In summary, our results elucidate how CIDE proteins are recruited to LDs and unveil a new mechanism by which PI(4)P regulates LD growth during adipogenesis and hepatic steatosis.

## MATERIALS AND METHODS

Key resources table

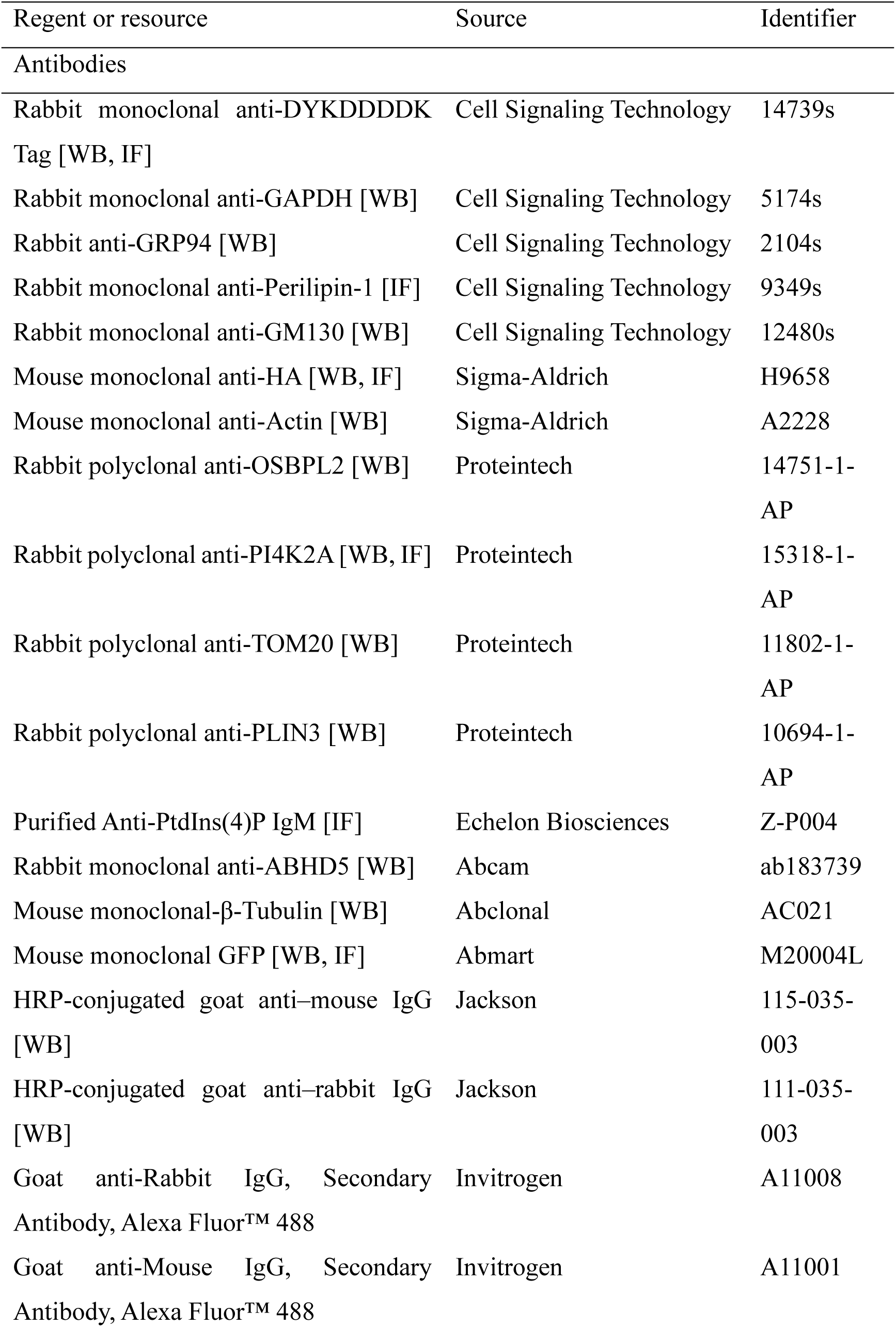
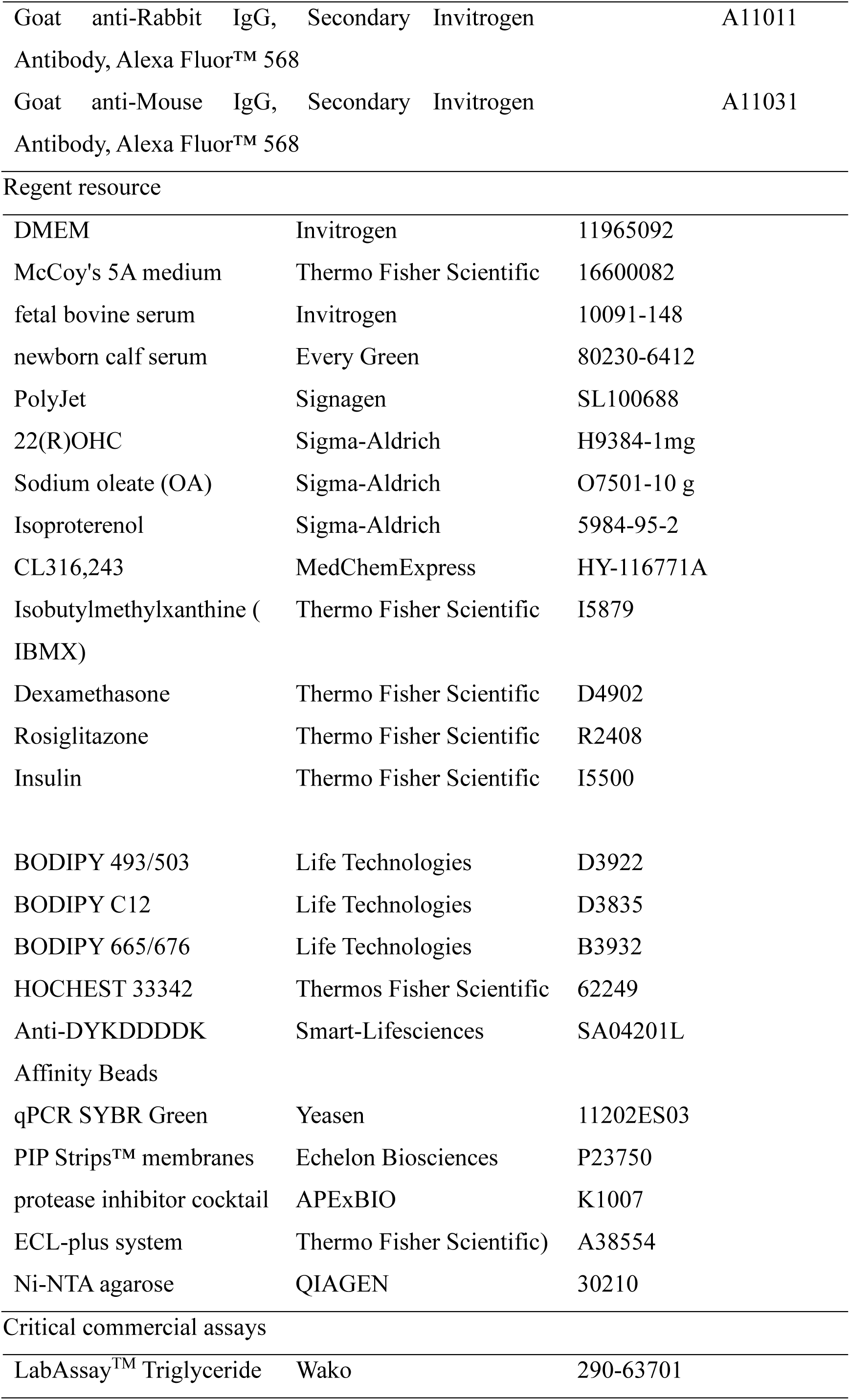
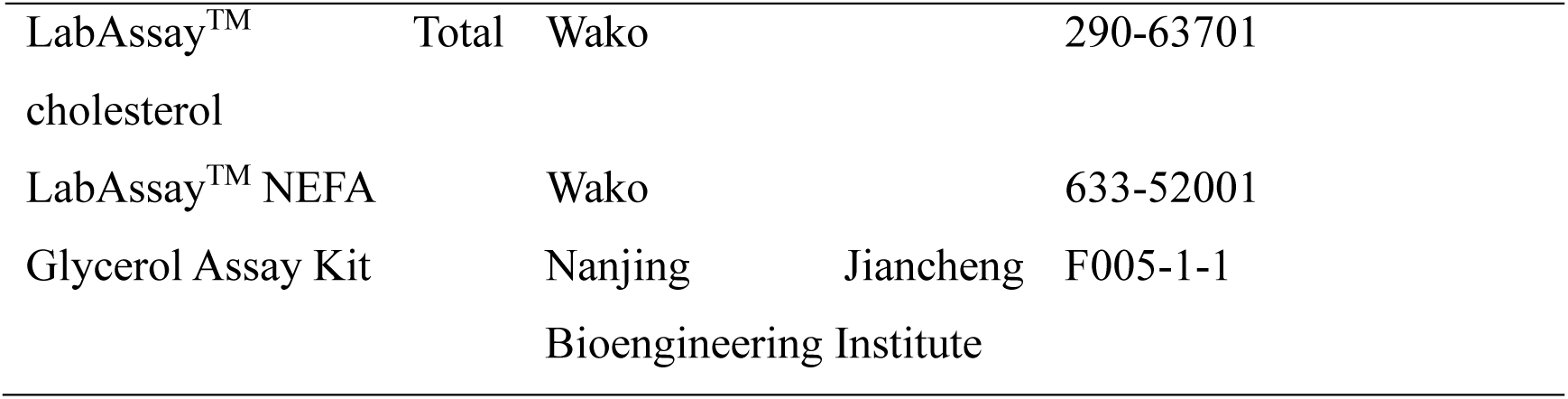

### Cell culture and transfection

HEK293T, HeLa, and Huh7 were cultured in Dulbecco’s modified Eagle’s Medium supplemented with 10% fetal bovine serum, 100 U/mL penicillin and 100 mg/mL streptomycin. U2-OS were cultured in McCoy’s 5A medium supplemented with 10% fetal bovine serum, 100 U/mL penicillin and 100 mg/mL streptomycin. 3T3-L1 pre-adipocytes were cultured in DMEM supplemented with 10% newborn calf serum, 100 U/mL penicillin and 100 mg/ml streptomycin. Cells were incubated at 37℃ in a humidified incubator containing 5% CO_2_. To promote the formation of lipid droplets (LDs), cells were treated with 50 μM or 200 μM OA conjugated to fatty acid-free BSA in water at a 6:1 molar ratio and cultured for 16 h. 3T3-L1 pre-adipocytes were induced to differentiate into mature adipocytes as described by the protocol of ATCC.

Plasmid DNAs were transfected into HEK293T, HeLa, Huh7, and U2-OS cells using PolyJet. Electroporation of plasmid DNAs into 3T3-L1 pre-adipocytes and 3T3-L1 mature adipocytes was performed using Amaxa Nucleofector II (Lonza) with a program A-033 according to the manufacturer’s instruction.

### Plasmid construction, mutagenesis, and RNA interference

Full-length cDNAs encoding various human proteins were amplified by PCR from the cDNAs of HeLa. cDNA encoding homo *ORP2*, *ORP5*, *CIDEA*, *CIDEB*, and *CIDEC* was cloned into pEGFP-C1 as previously described ^9, 27^. Point mutations or domain deletions of *ORP2* (W77A, I79A, HH/AA, ΔFFAT, ΔELSK, and ΔHelix), *ORP5* (W372A, L374A, HH/AA, and ΔTM), and *PI4K2A* (K152A and D308A) were introduced or generated by PCR-based site-directed mutagenesis/deletions. cDNA encoding mus *Plin1*, *Plin2*, *Plin3*, *Abhd5*, *Dgat2*, *Rab18/homo PLIN1*, *PLIN2*, *PLIN3*, *ABHD5*, *DGAT2*, *RAB18* was cloned into pEGFP-C1 or pEGFP-N1vectors. cDNA encoding homo *ORPs* family members was cloned into pEGFP-C1 as previously described^27^. The siRNAs used for knockdown of *ORP2*, *ORP5*, *PI4KA*, *PI4KB*, *PI4K2A*, and *PI4K2B* are listed in Table S1. All siRNAs were synthesized by GenePharma. The primer sequences used for real-time PCR analysis are listed in Table S2.

### Lipid strip assay

Recombinant wild-type CIDEC and mutants were expressed in BL21 (DE3) cells. Protein expression was induced by adding 1 mM isopropyl beta-d-thiogalactopyranoside (IPTG) overnight at 18℃. Cells were collected and lysed by lysis buffer (50 mM Tris-HCl, pH 7.4, 300 mM NaCl, 1 mM TCEP). Centrifuge-cleared lysate was applied to Ni-NTA agarose, washed with lysis buffer containing 20 mM imidazole. Purified CIDEC proteins were collected and probed with Membrane Lipid Strips according to the manufacturer’s instructions.

### High-content screening system (HCS)

HeLa cells were implanted in imaging-compatible 96-well plates, transfected with an indicated siRNA in every well using RNAiMAX, and fixed 48 h after transfection. Then, Nuclei were stained with HOCHEST 33342, LDs were stained with BODIPY 493/503, and the multi-layer images of intracellular LDs were captured by a high content imaging platform (Opera Phenix, Perkin Elmer). Next, multi-layer images were analyzed using the Harmony software to produce a 3D image, and nucleus and intracellular LDs were identified to obtain the parameters for the number of nuclei and the number and volume of LDs. Two quantitative indicators were obtained by calculation: LD number/cell and LD volume/LD.

### Immunofluorescence

Cells were rinsed twice in PBS, fixed with 4% PFA (at 4°C for 6-8 h or at room temperature for 15 min), permeabilized and blocked with 1%Triton X-100 and 1.5% BSA in PBS for 30 min, followed by incubation with primary antibody for 2 h, washed three times with PBS, and incubated with fluorescently labeled secondary antibody 1:500 dilution for 1 h followed by BODIPY dyes if required. Images of cells with LDs and nuclei were acquired under a confocal laser scanning microscope (Zeiss LSM 880) with an optical sectioning by Apotome module and a 63× oil immersion objective.

### Image Analysis

The brightness and contrast of fluorescent confocal images, as well as further analysis of images (e.g., amplifying a certain region), were adjusted in parallel using ImageJ (NIH). The Pearson’s correlation coefficient of two fluorescence signals was calculated by the ZEISS LSM 880 blue software. The LD volume or area and LD number in a cell,H&E and immunofluorescence image were measured and quantified using ImageJ (NIH) or IMARIS 9.5 (Oxford instruments).

### Lentivirus preparation and infection

The preparation of lentivirus was performed in HEK 293T cells. For lentiviral packaging, an indicated gene on pCDH-puro or a shRNA on pLKO.1 was co-transfected with psPAX2 and pMD2.G into the cells. For infection, HeLa cells were infected at the confluence of 40-60% with lentiviruses. After infection, the HeLa cells were antibiotically selected against 1 μg/mL puromycin for at least 48 h.

### LD isolation

Cells treated with OA in medium overnight were washed twice with PBS, collected by scraping into PBS, and pelleted by centrifugation at 1,000 *g*, 4°C for 10 min. Cell pellets were resuspended in TES buffer (20 mM Tris, pH 7.4, 1 mM EDTA, and 250 mM sucrose) and dounced for 50 times. The lysate was centrifuged at 500 *g*, 4°C for 10 min. The postnuclear supernatant was then transferred into SW 41 Ti tubes and hypotonic buffer (10 mM HEPES, 1 mM EGTA and 25 mM KCl, pH 7.8.) was loaded on top of the postnuclear supernatant, and the gradient was subjected to sequential ultracentrifugation (OptimaTM L-100 XP Ultracentrifuge) at 38500 rpm at 4 °C for 1 h and 40 mins. After each ultracentrifugation step, LDs floating on the upper layer of the centrifuge tubes were carefully collected and washed with hypotonic buffer three times.

### Animals and AAV injection

The *Osbpl2*-specific, *Osbpl5* ORF-specific, and *Pi4k2a*-specific shRNAs were inserted in the AAV9-pCAG-EGFP vector (Addgene plasmid no. 51502) or AAV8-TBG-EGFP vector (Addgene plasmid no. 105535). Then 10^12^ viral genomes per mouse were injected through subcutaneously in the inguinal white adipose tissue of wild-type mice or intravenously in the liver of *ob/ob* mice. To promote lipolysis, each wild-type mouse received an intraperitoneal injection of CL316,243 at 1 mg/kg/d for 5 days or fasting for 20 h. Mice were housed in the animal facility with 12-h light/ 12-h dark cycles, the temperature at 22°C–23°C and 20-60% humidity, with free access to standard chow and water. All animal procedures were approved by the Institutional Ethics Committee of Fudan University, under a permit of animal use in the Center of Experimental Animal at Shanghai. The permit followed the Experimental Animal Regulations set by the National Science and Technology Commission, China.

### FRAP-based lipid exchange rate assay

FRAP-based lipid exchange rate assay was performed as described previously^13, 39^. Briefly, CIDEC HeLa cells were incubated with 200 µM OA and 1 µg/ ml BODIPY C_12_ for 16 h and transferred to fresh medium 1 h before FRAP experiments. Live cells were visualized under a confocal microscope (A1Rsi, Nikon) using a 100× oil-immersion objective. At least 70% of LD total area was photobleached for 1 s at 561-nm laser, followed by time-lapse scanning with a 3 s interval. The same photobleaching process was repeated three times. Mean optical intensities (MOIs) within LD core regions were measured simultaneously. We further calculated the rate of lipid exchange between LD pairs randomly we selected as previously^9^.

### Co-immunoprecipitation and Western blot analysis

Co-immunoprecipitation and Western blot analyses were performed according to the previous procedure^40^. Cells transfected with indicated plasmids were washed with PBS and cellular proteins were extracted with IP buffer (1% Sodium Deoxycholate, 1% NP-40, 5 mM EGTA, 5 mM EDTA, protease inhibitor cocktail by sonication. Flag agarose beads were used for immunoprecipitation. Sample were subjected to Western blot analysis with the indicated antibodies. The blots were detected using secondary HRP-conjugated antibodies under the ECL-plus system.

### Statistical Analysis

Data represents the mean ± SD or mean ± SEM of at least three independent experiments as indicated in figure legends. P values lower than 0.05 were considered significant. Significance for two-group comparison was established using a two-tailed Student’s *t* test. Statistical difference was shown as \**P* < 0.05, \*\**P* < 0.01, and \*\*\**P* < 0.001. ^NS^*P* > 0.05 represents no significant difference.

## ACKNOWLEDGEMENT

This work was supported by grants from the National Natural Science Foundation of China (92357308, 92357302, and 32270724 to F.J.C. and P.L.), the National Key R&D Program of China (2018YFA0800301 to F.J.C.), and the High-Level Medicine Foundation of Shanghai Government (to P. L.). This work is also supported by Shanghai Basic Research Field Project “Science and Technology Innovation Action Plan” (Grant No. 21JC1400400) and Shanghai Municipal Science and Technology Major Project (Grant No. 2017SHZDZX01). H.Y. was supported by an Investigator Grant (2009852 from the National Health and Medical Research Council (NHMRC) of Australia, and by start-up funding from the University of Texas Health Science Center at Houston.

The authors declare no competing financial interests.

## Author contributions

J. Wu, M. Gao and X. Du performed experiments. All authors analyzed data. J. Wu, F.J. Chen, P. Li and H. Yang designed research. H. Yang wrote the manuscript with input from all other authors.

## Online Supplemental Material

**Figure S1.**
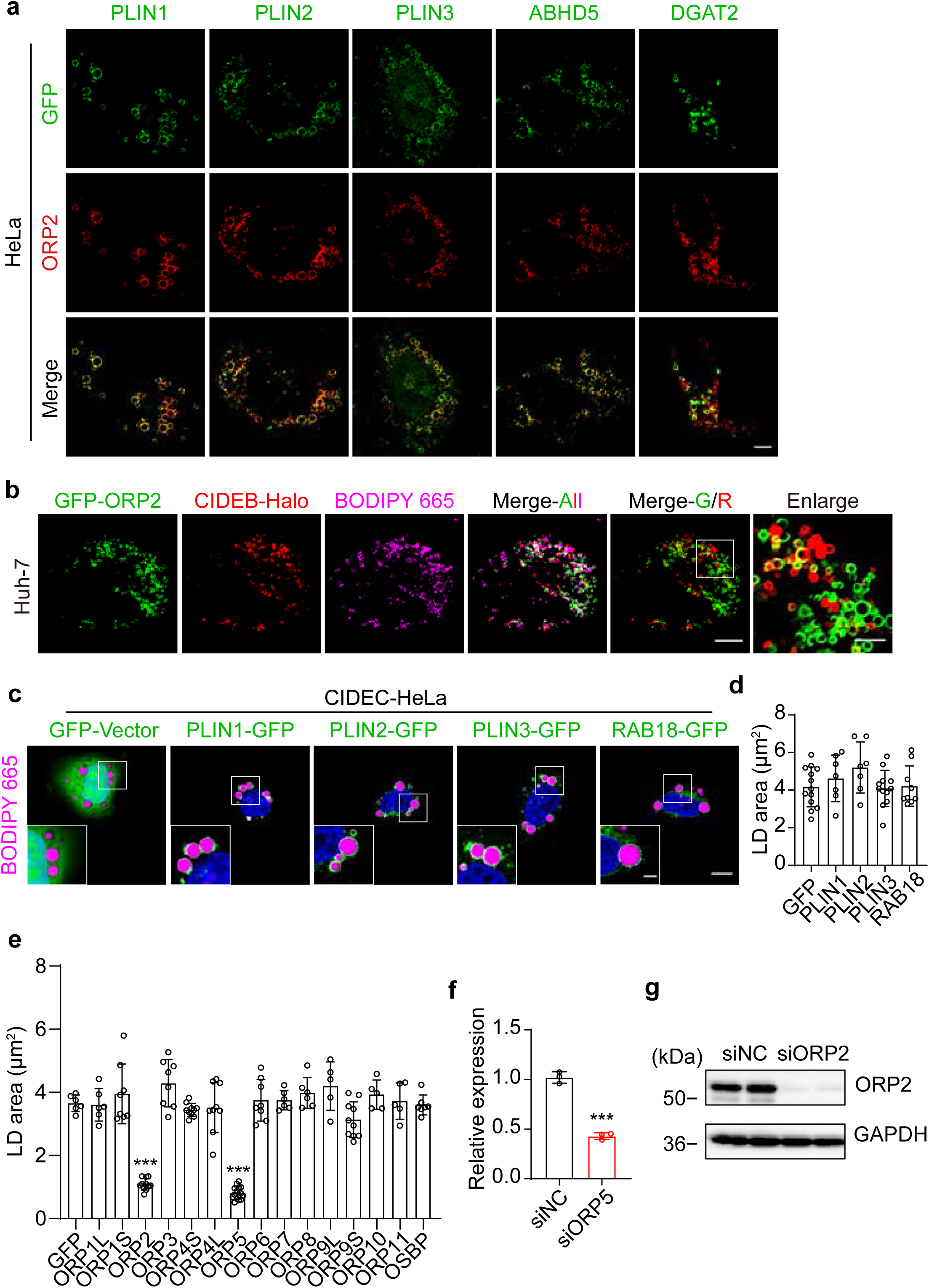
ORP2/5 affect the localization and function of CIDE proteins but not other LD-associated proteins. a. Representative confocal images showing localizations of ORP2 and LD targeting proteins such as PLIN1, PLIN2, PLIN3, ABHD5, or DGAT2 in wild-type HeLa cells. Green, PLIN1, PLIN2, PLIN3, ABHD5, and DGAT2. Red, ORP2. Scale bar, 10 µm. b. Representative confocal images showing the localization of ORP2 and CIDEB in Huh7 cells treated with 200 μM OA for 16 h. Green, GFP-ORP2. Red, CIDEC-Halo. Magenta, LDs (BODIPY 665). Scale bars, 10 µm; (Enlarged) 3 µm. c. Representative confocal images showing the LD morphology in CIDEC-Hela cells overexpressing LD targeting proteins such as PLIN1, PLIN2, PLIN3, or RAB18. Green, GFP, PLIN1, PLIN2, PLIN3, or RAB18 proteins. Magenta, LDs (BODIPY 665). Blue, nuclei (DAPI). Scale bars,10 µm; (Enlarged) 3 µm. d. Histogram showing the quantification of average LD area in (c). (n=7-15) e. Histogram showing the quantification of average LD area in CIDEC-HeLa cells individually overexpressing GFP-tagged ORPs as indicated. (n=5-15) f. Histogram showing a reduction in *ORP5* mRNA expression in *ORP5*-knockdown wild-type HeLa cells after treatment with *ORP5* siRNA (siORP5), compared to that treated with scramble siRNA (siNC). Three independent experiments. g. Western blotting showing the expression level of ORP2 in wild-type HeLa cells after treatment with scrambled siRNA (siNC) or *ORP2* siRNA (siORP2). GAPDH served as a loading control.

**Figure S2.**
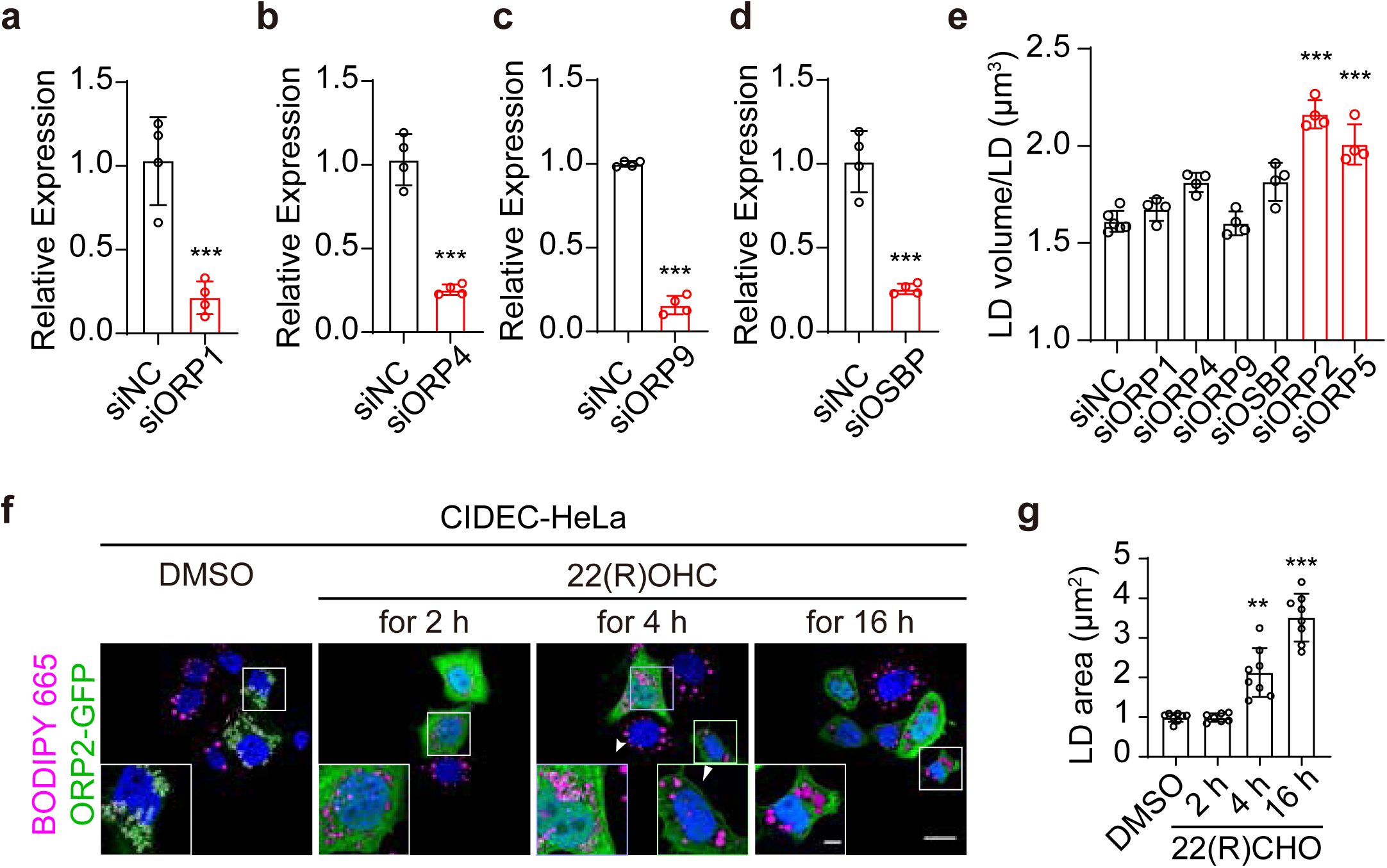
Localization of ORP2/5 to LDs is necessary for their impact on LD size. a-d. Histogram showing reductions in indicated mRNA expression in wild-type HeLa cells after treatment with *ORP1* siRNA (siORP1, a), *ORP4* siRNA (siORP4, b), *ORP9* siRNA (siORP9, c), and *OSBP* siRNA (siOSBP, d), respectively, compared to that treated with scramble siRNA (siNC). e. Histogram showing the quantification of LD volume/LD in CIDEC-HeLa cells after treatment with scrambled siRNA (siNC), *ORP1* siRNA (siORP1), *ORP4* siRNA (siORP4), *ORP9* siRNA (siORP9), *OSBP* siRNA (siOSBP), *ORP2* siRNA (siORP2), or *ORP5* siRNA (siORP5) analyzed by high-content screening system for four independent experiments. f. Representative confocal images showing the LD morphology in CIDEC-HeLa cells overexpressing GFP-ORP2 followed by treatment with 22(R)OHC for 2, 4, and 16 h, compared to that treated with DMSO. Green, GFP-ORP2. Magenta, LDs (BODIPY 665). Blue, nuclei (DAPI). Scale bars, 20 µm; (Enlarged) 3 µm. g. Histogram showing the quantification of LD area in CIDEC-HeLa cells in (f). (n=8-10)

**Figure S3.**
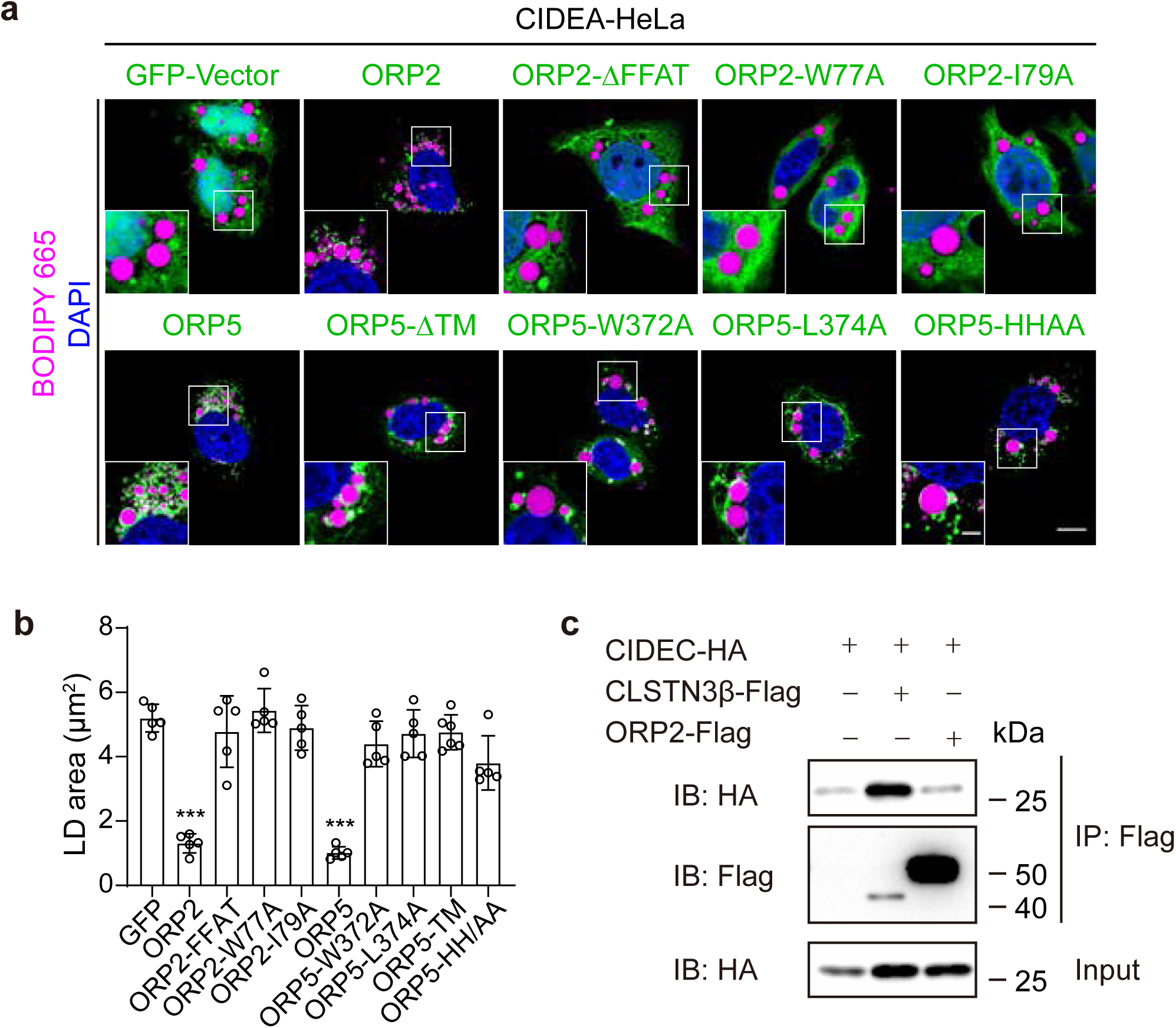
ORP2/5 inhibited CIDEA function. a. Representative confocal images showing the LD morphology in CIDEA-HeLa cells overexpressing indicated proteins. Green, GFP, wild-type ORP2, OPR2 mutants, wild-type ORP5, and ORP5 mutants. Magenta, LDs (BODIPY 665). Blue, nuclei (DAPI). Scale bars, 10 µm; (Enlarged) 3 µm. b. Histogram showing the quantification of LD size in CIDEA-HeLa cells in (a). (n=5-10) c. Western blotting assay showing co-immunoprecipitation of ORP2-Flag or CLSTN3β-Flag with CIDEC-HA expressed in HEK293T cells.

**Figure S4.**
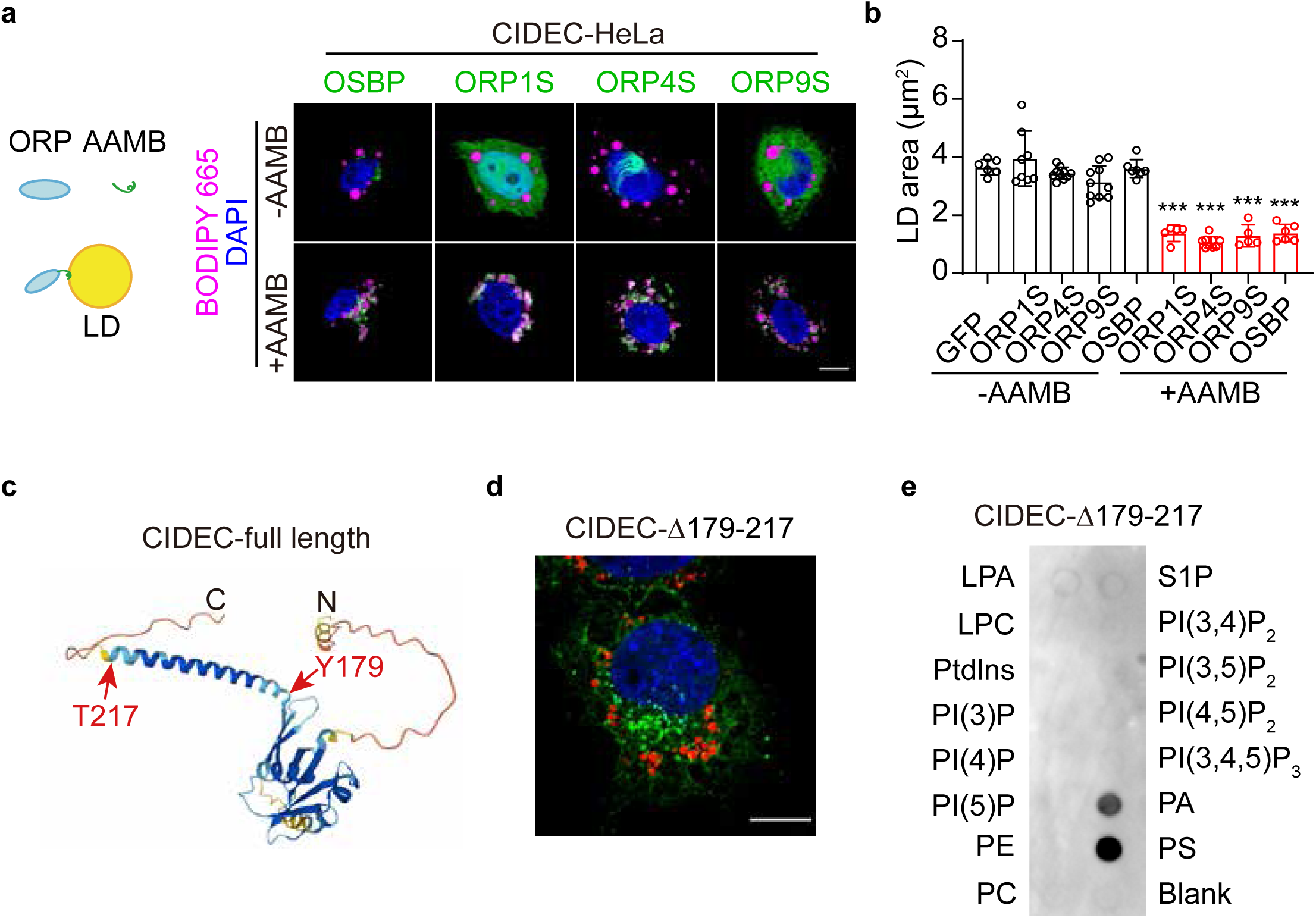
a. Left: Schematic diagram showing ORP proteins fused with the LD targeting motif of AAMB can be localized to the LD surface. Right: Representative confocal images showing the localizations of OSBP-GFP, ORP1S-GFP, ORP4S-GFP, ORP9S-GFP with or without AAMB in CIDEC-HeLa cells treated with 200 μM OA. Green, OSBP and ORPs. Magenta, LDs (BODIPY 665). Blue, nuclei (DAPI). Scale bars, 10 µm. b. Histogram showing the quantification of average LD area in (a). (n=5-10) c. The predicted structure of CIDEC protein by Alphafold. d. Confocal images showing the localization of CIDEC-Δ179-217 in HeLa. e. Lipid strip assay showing the affinity of CIDEC-Δ179-217 for phospholipids.

**Figure S5.**
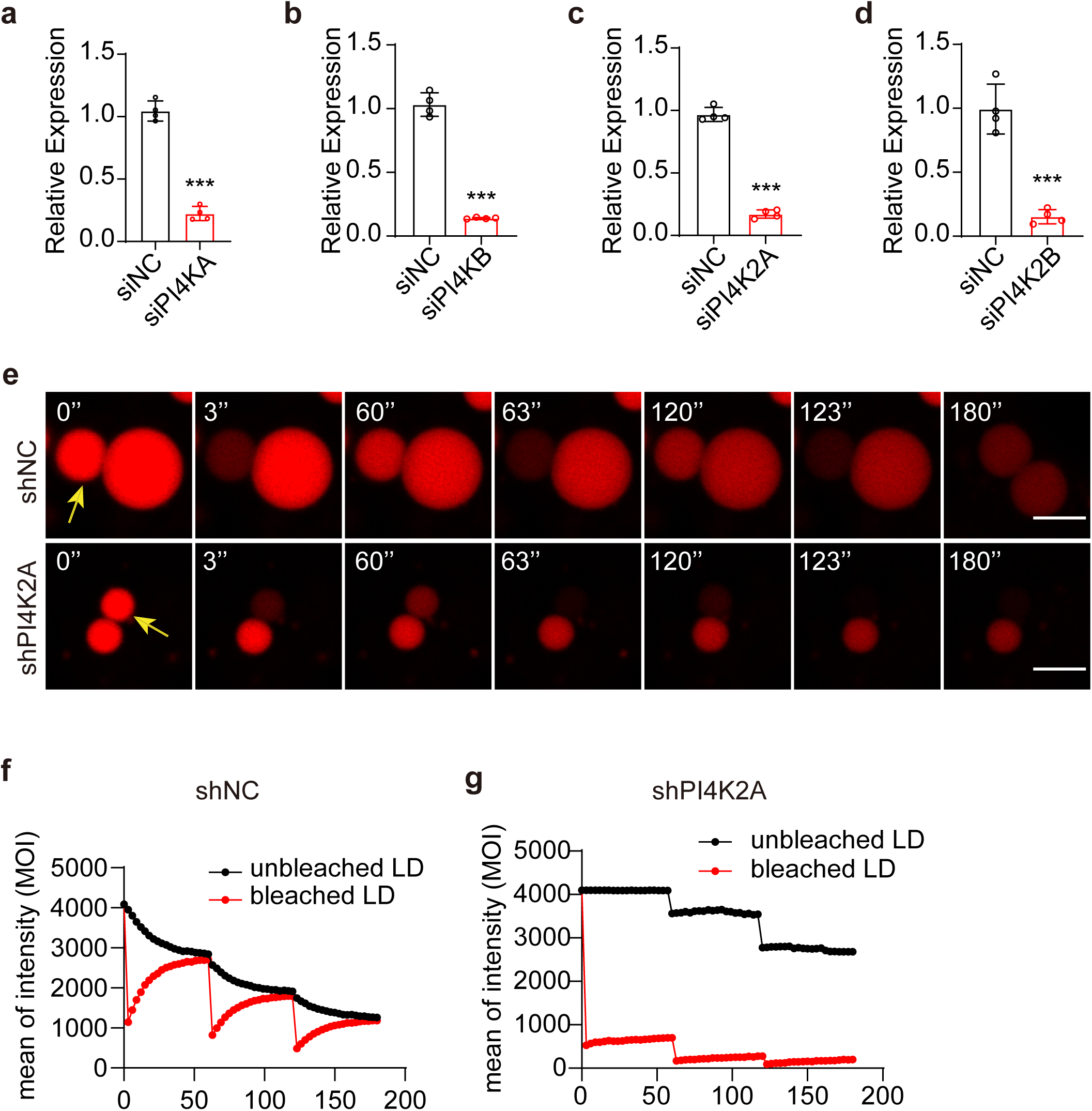
PI4K2A regulates CIDEC-mediated lipid exchange rates. a-d. Histograms showing reductions in indicated mRNA expression in wild-type HeLa cells after treatment with *PI4KA* siRNA (siPI4KA, a), *PI4KB* siRNA (siPI4KB, b), *PI4K2A* siRNA (siPI4K2A, c), and *PI4K2B* siRNA (siPI4K2B, d), respectively, compared to that treated with scrambled siRNA (siNC). Four independent experiments. e-g. Representative images (e) and mean optical intensity traces (f) and (g) of FRAP-based lipid exchange rate assay for one pair of LDs in CIDEC-HeLa cells infected with lentiviral scrambled shRNA (shNC) or *PI4K2A* shRNA (shPI4K2A). LDs were pre-stained with BODIPY C12. Scale bars, 3 µm.

**Figure S6.**
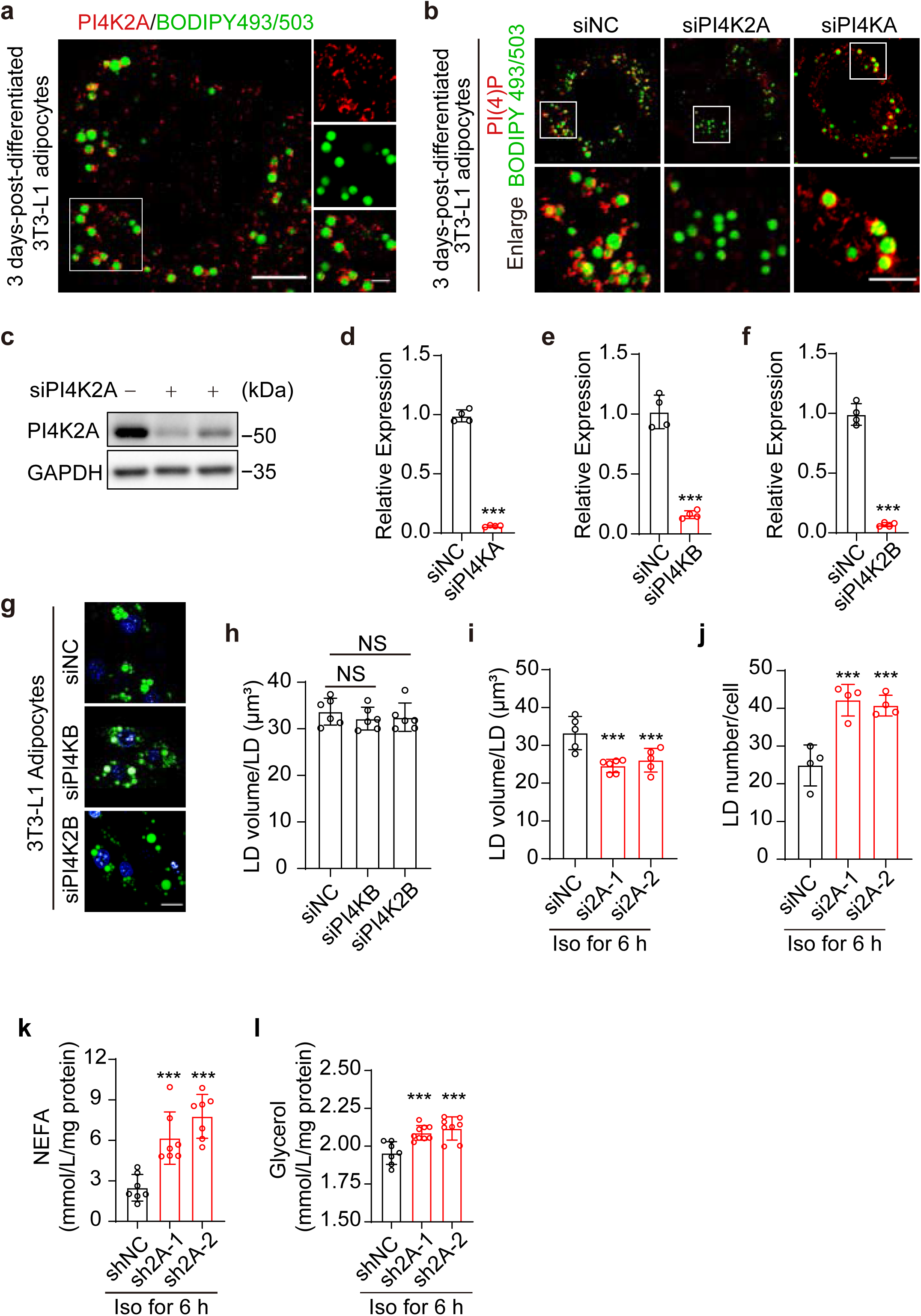
PI4K2A regulates LD morphology and lipolysis in 3T3-L1 adipocytes. a. Representative confocal images showing endogenous PI4K2A detected by specific antisera against PI4K2A in 3 days-post-differentiated 3T3-L1 adipocytes. Green, LDs (BODIPY 493/503). Red, PI4K2A. Scale bars, 10 µm; (Enlarged) 3 µm. b. Representative confocal images showing PI(4)P detected by specific against PI(4)P in 3 days-post-differentiated 3T3-L1 adipocytes after treatment with scrambled siRNA (siNC), *PI4K2A* siRNA (siPI4K2A), or *PI4KA* siRNA (siPI4KA). Green, LDs (BODIPY 493/503). Red, PI(4)P. Scale bars, 10 µm; (Enlarged) 3 µm. c. Western blotting showing the expression level of PI4K2A in 3T3-L1 pre-adipocytes after treatment with scrambled siRNA (siNC) or *PI4K2A* siRNA (siPI4K2A). GAPDH served as a loading control. d-f. Histograms showing mRNA expression of indicated genes in 3T3-L1 pre-adipocytes after treatment with *PI4KA* siRNA (siPI4KA, d), *PI4KB* siRNA (siPI4KB, e), and *PI4K2A* siRNA (siPI4K2A, f), respectively, compared to that treated with scrambled siRNA (siNC). Four independent experiments. g. Representative confocal images showing the LD morphology in 5 days-post-differentiated 3T3-L1 adipocytes after treatment with scrambled siRNA (siNC), *PI4KB* siRNA (siPI4KB), or *PI4K2B* siRNA (siPI4K2B), respectively. Green, LDs (BODIPY 493/503). Blue, nuclei (DAPI). Scale bar, 10 µm. h. Histogram showing the quantification of LD volume/LD in 3T3-L1 adipocytes as in (g) analyzed by high-content screening system. (n=6) i and j. Histograms showing the quantification of LD volume/LD (i) and LD number/cell (j) in 3T3-L1 adipocytes after treatment with *PI4K2A* siRNA at 4 days of post-differentiated adipocytes compared to that treated with scrambled siRNA (siNC), followed by treatment with 1 μM isoproterenol (Iso) for 6 h analyzed by high-content screening system. (n=4-6) k and l. Histograms showing the measurements of non-esterified fatty acids (NEFA) (k) and glycerol release (l) of 3T3-L1 adipocytes infected with lentiviral GFP-tagged scrambled shRNA (shNC) or *PI4K2A* shRNA (shPI4K2A) at 4 days-post-differentiated adipocytes followed by treatment with 1 μM Iso for 6 h. (n=5-10)

**Figure S7.**
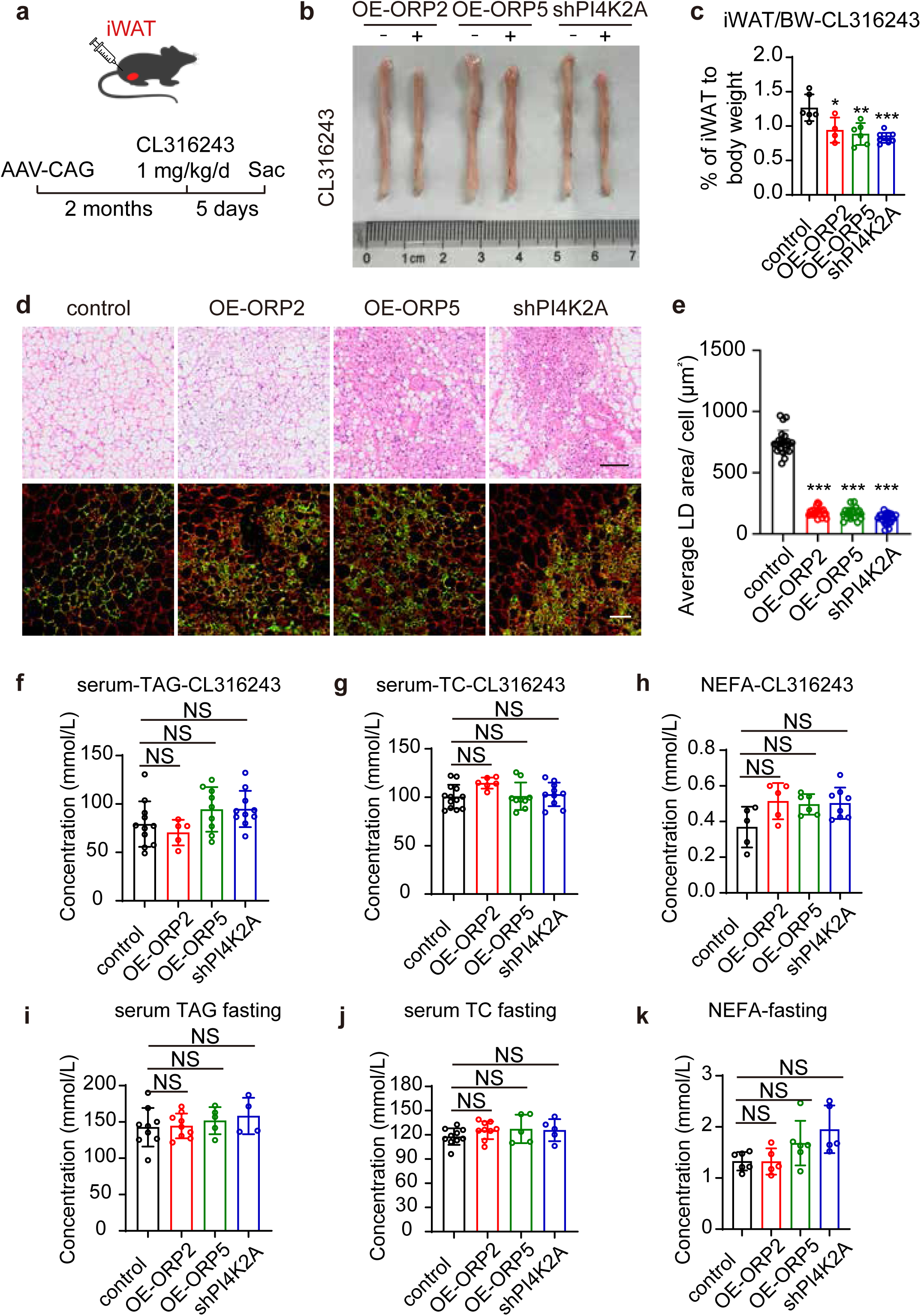
ORP2/5 overexpression or *PI4K2A* knockdown promotes lipolysis in the adipose tissues of wild-type mice after treatment with CL316,243. a. Schematic diagram showing that 8-week-old male mice were injected *in situ* with AAV-GFP, AAV-GFP-ORP2, AAV-GFP-ORP5, or AAV-GFP-shPI4K2A into the iWAT for 2 months followed by treatment with CL316,243 (1 mg/kg/d) for 5 days. n=4-9 mice. Sac, sacrifice. b. Representative images showing the iWAT of mice in (a) treated with CL316,243 for 5 days. c. Histogram showing the ratio of iWAT mass to body weight (iWAT/BW) of mice after treatment with CL316,243 for 5 days. d. H&E staining and immunofluorescence (red, Perilipin1, green, GFP) images showing the iWAT of mice in (a). Scale bars, 100 µm (H&E) and 50 µm (IF). e. Histogram showing the quantification of average LD area of iWAT in (d). f-h. Histograms showing the quantification of triglycerides (TG) (f), total cholesterol (TC) (g), non-esterified fatty acids (NEFA) (h) in the serum of mice in (a). (n=5-9) i-k. Histograms showing the quantification of triglycerides (TG) (i), total cholesterol (TC) (j), non-esterified fatty acids (NEFA) (k) in the serum of mice administered by *in situ* injecting with AAV-GFP, AAV-GFP-ORP2, AAV-GFP-ORP5, or AAV-GFP-shPI4K2A into the iWAT followed by fasting for 20 h.

**Figure S8.**
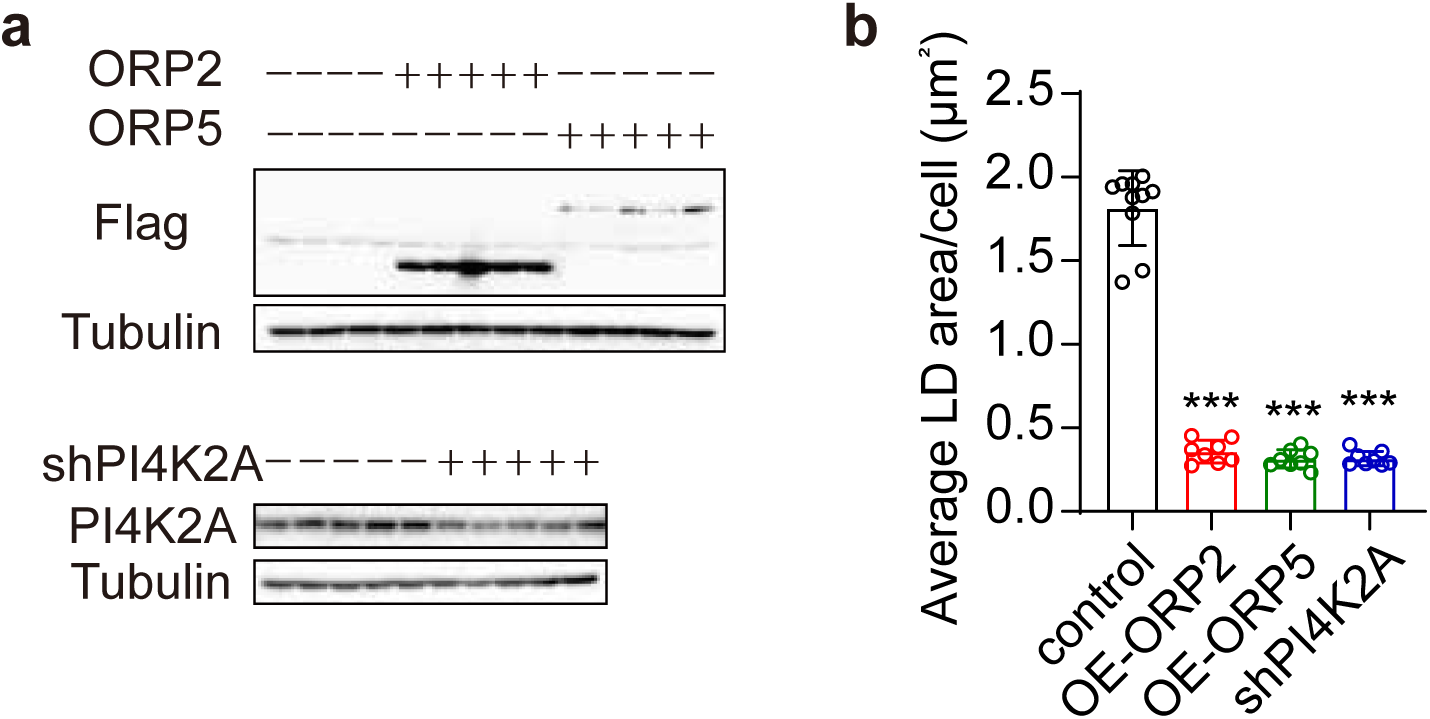
ORP2/5 overexpression or *PI4K2A* knockdown in liver of ob/ob mice. a. Western blotting showing the level of indicated protein in the liver of *ob/ob* mice. b. Histogram showing the quantification of average LD area of liver as in Fig. 7j-7k.

**Figure S9.**
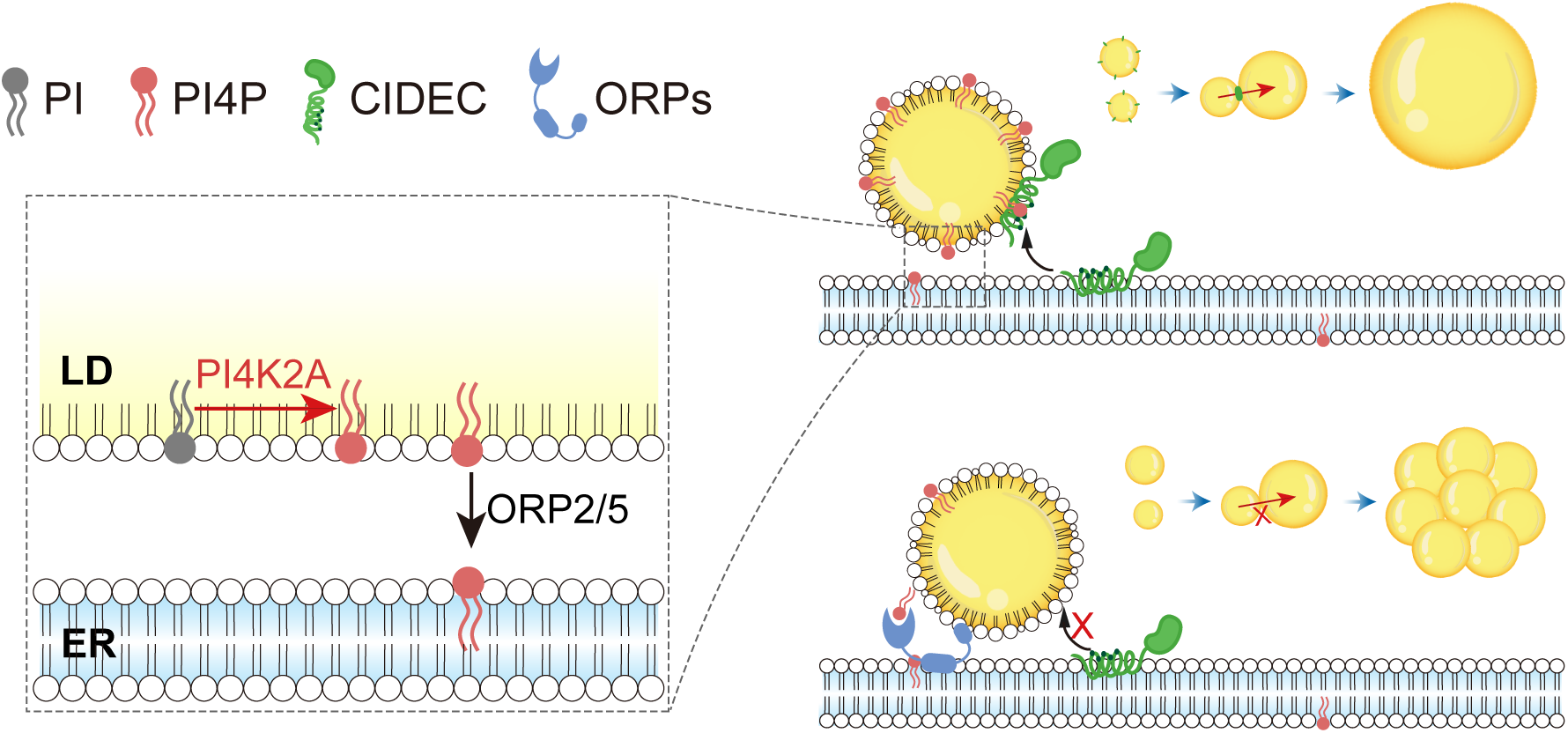
PI(4)P is present on the surface of lipid droplets and controls CIDE protein function. PI(4)P can directly bind CIDEC and recruit it to LDs. Subsequently, CIDEC promotes the formation of unilocular white adipocytes. The level of PI(4)P on LDs is regulated by PI4K2A and ORP2/5. On the one hand, PI(4)P is synthesized by PI4K2A on the LD. On the other hand, PI(4)P is consumed by ORP2/5 to drive the delivery of cholesterol/phosphatidylserine to LDs. Without enough LD surface PI(4)P, a multilocular morphology of adipocytes is manifested because CIDE-mediated LD growth is suppressed.

**Table S1.**
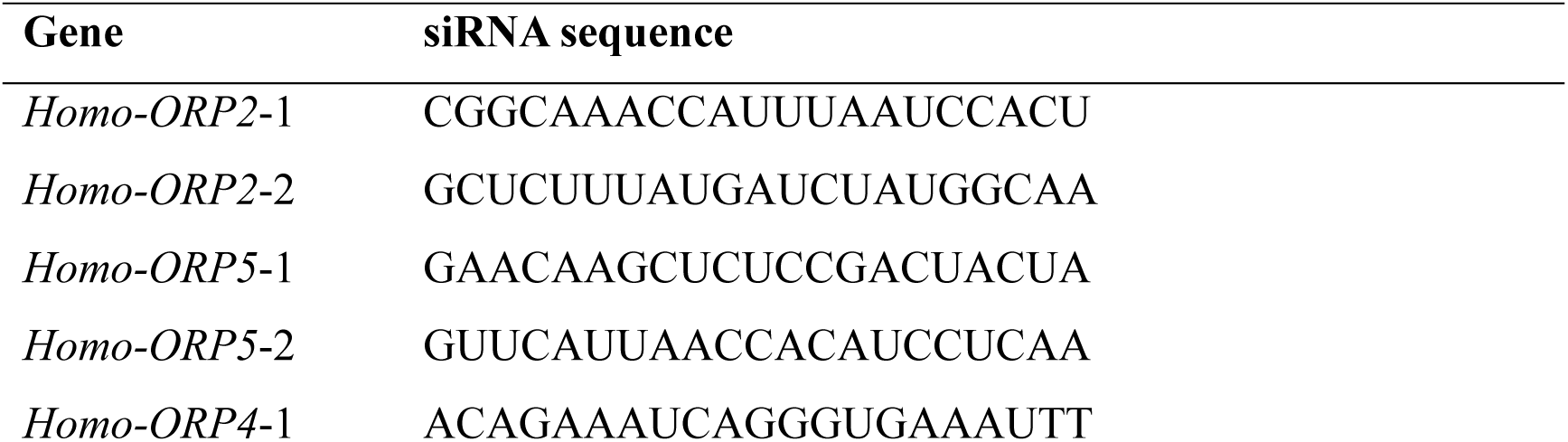
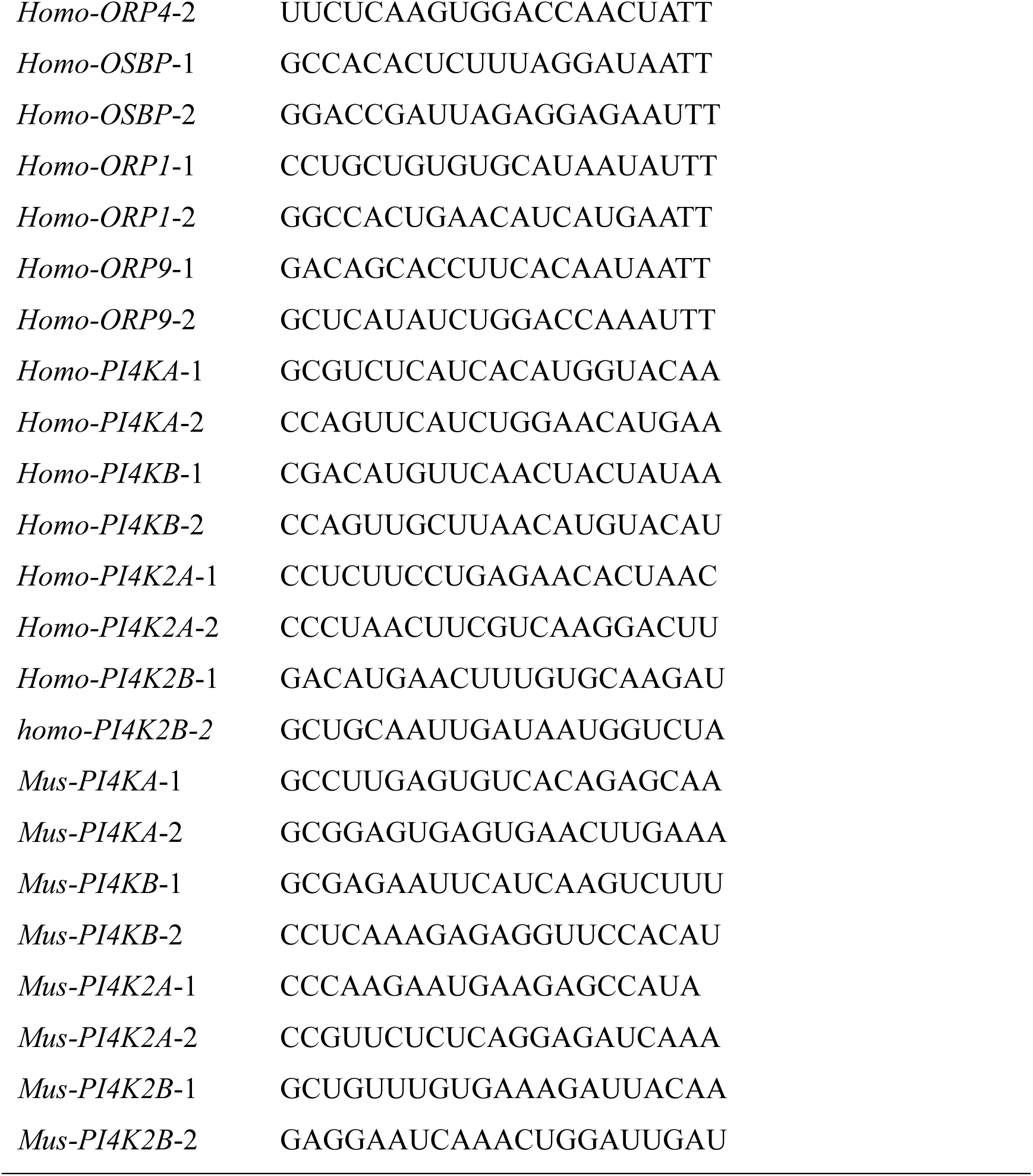
List of siRNA sequence used in this study.

**Table S2.**
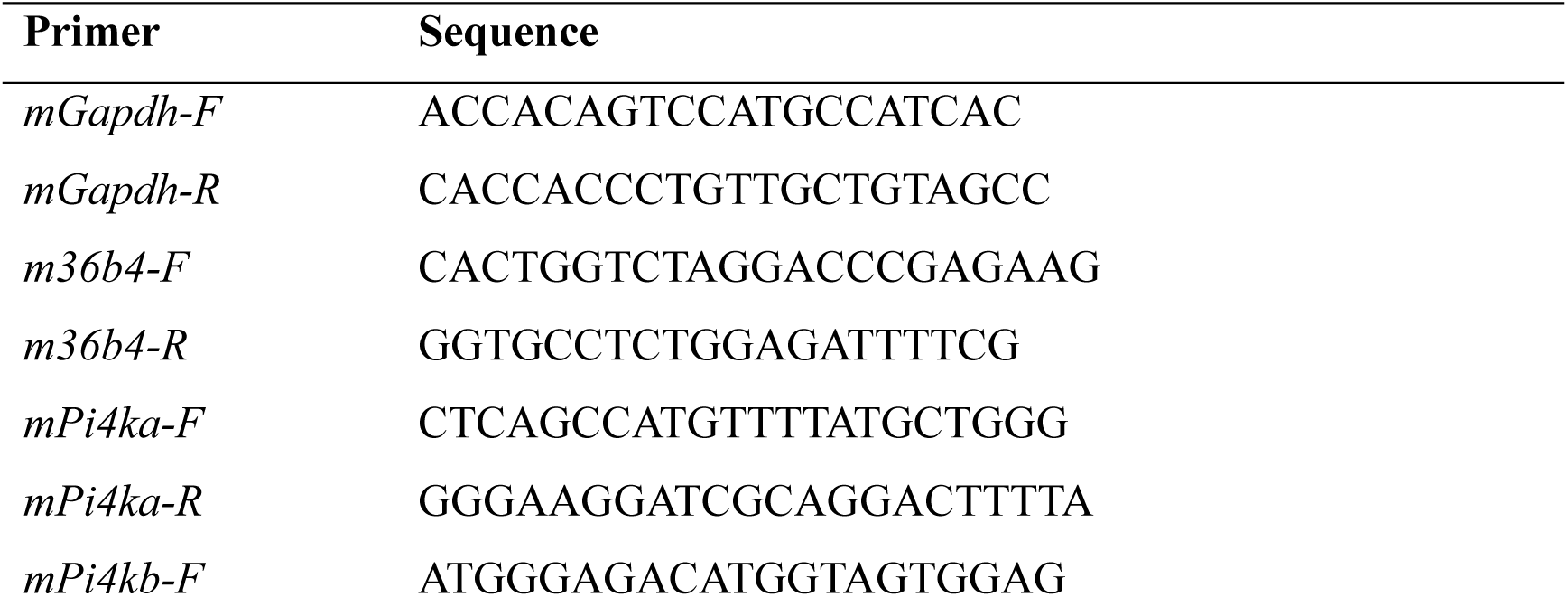
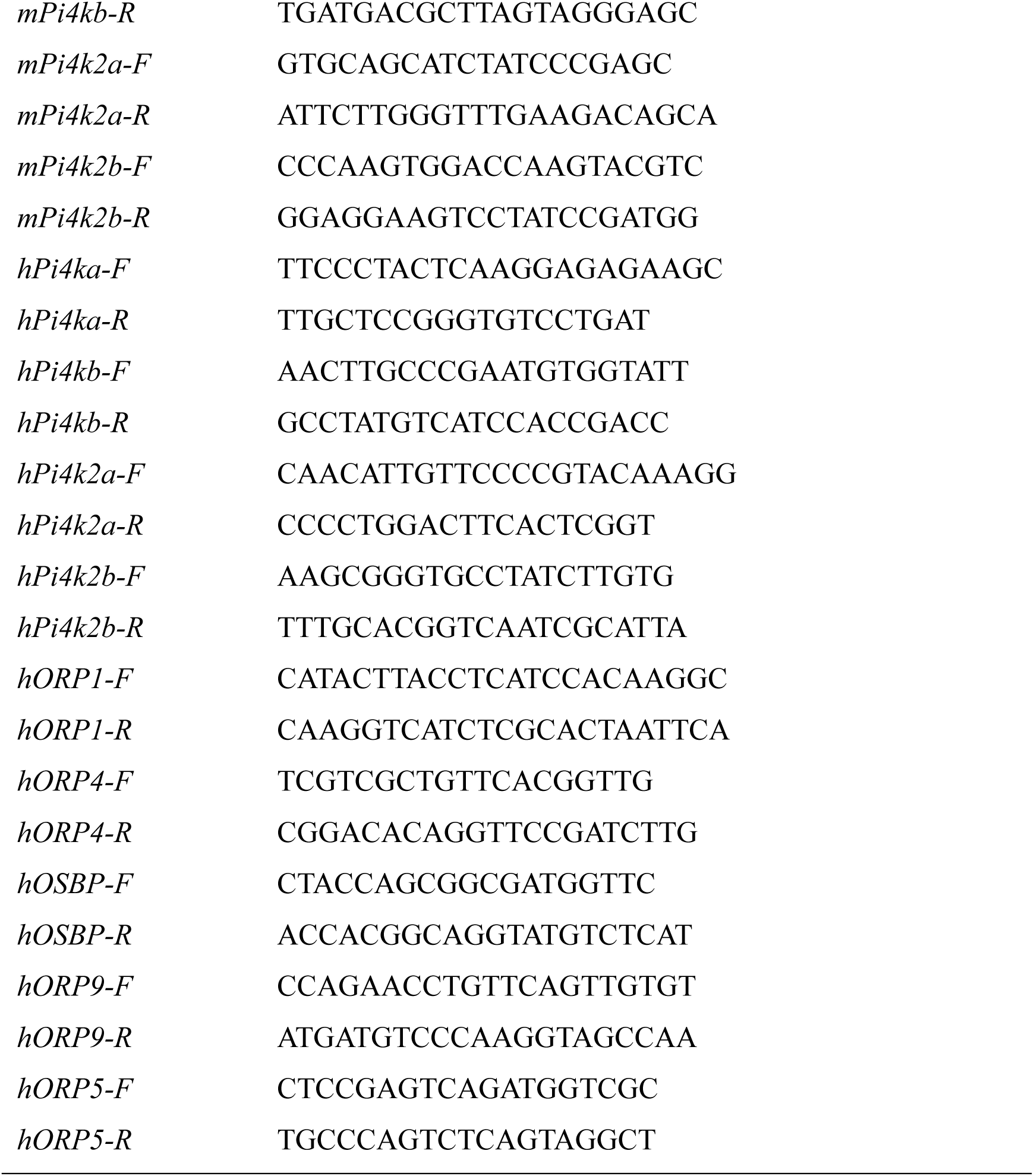
List of the primers used in this study.

